# T cell receptor-ligand affinity quantitatively tunes transcriptome remodelling *in vivo* inversely regulating cell division and interferon response

**DOI:** 10.64898/2026.06.24.734351

**Authors:** Veera Panova, Andrew Muir, Erick Armingol, Oliver Burton, Jessica Quantrill, David A Lewis, Simon J Cook, Martin Turner, Adrian Liston, Roser Vento-Tormo, Arianne C Richard

**Affiliations:** Immunology Programme, Babraham Institute, Babraham Research Campus, Cambridge, UK; Wellcome Sanger Institute, Wellcome Genome Campus, Hinxton, UK; Department of Pathology, University of Cambridge, Cambridge, UK; Department of Infectious Disease, Imperial College London, London, UK; The Francis Crick Institute, London, UK; Signalling Programme, Babraham Institute, Babraham Research Campus, Cambridge, UK

## Abstract

The strength of T cell receptor (TCR) engagement by antigenic ligands governs naïve T cell activation, expansion and differentiation. However, the molecular changes underpinning this process *in vivo* remain incompletely understood. To address this, we coupled an influenza infection model of varied TCR-ligand affinity with high-dimensional protein and RNA measurements. As previously reported, high affinity stimulation drove greater expansion, with disproportionately abundant effector subsets. Experiments using a KLRG1-fate reporter showed that differentiation state biases affected cells from both direct and indirect memory differentiation trajectories. Early post-infection, these biases manifested in differential metabolic and proliferative activities. Examination of T cells showing signs of initial priming *in vivo* revealed that TCR-ligand affinity primarily changed the magnitude of TCR-induced transcriptome remodelling, not the genes involved. This quantitative tuning drove affinity-dependent amplification of ribosome biogenesis and suppression of interferon response genes before mitogenic diversion, and was associated with elevated TCR-induced transcription factor activity. Together, these data demonstrate the *in vivo* underpinnings of affinity-dependent responses and reveal how accumulation of TCR-induced signalling outputs translates binding properties into appropriate differentiation outcomes.

## INTRODUCTION

CD8^+^ T cells play a central role in combating intracellular pathogens and eliminating cancerous cells (1,2). Naïve T cells become activated when peptide-MHC ligand complexes bind T cell receptors (TCRs) (3) alongside co-stimulatory signals (4,5), initiating gene expression and metabolic changes that enable proliferation and differentiation into functional subsets. In the days following antigen stimulation, activated CD8^+^ T cells separate into short-lived effector and memory precursor populations (6–10), whose unique expression profiles are accompanied by differences in proliferation rate (11–13). However, fates remain plastic, such that effector-phenotype lineages can later divert to memory states (14,15). While individual naïve T cells stochastically give rise to progeny with heterogeneous differentiation fates, proportional distributions of effector and memory subsets are relatively stable at the population level (16–20) and can be skewed by variations in antigenic or other immune environment signals (21).

Previous work has shown that the strength of TCR-antigen interactions controls the magnitude of subsequent expansion (5,22–24). While very weak ligands have limited ability to trigger TCR signalling cascades or proliferation, there is a substantial range of TCR-ligand affinities in which activation can be achieved (25–27). *In vitro* studies have found both graded and on-off behaviours among TCR-induced signalling and gene expression changes that are tuned by antigen affinity (28–32), as well as a probabilistic mechanism in which TCR-ligand affinity determines the rate with which naïve T cells initiate common signalling, transcriptional and proliferative programs (33–35).

However, the *in vivo* environment of an active lymph node is complex, including spatial variation in cellular and soluble stimuli, as well as repeated antigen encounters (36), that provide further opportunities for ligand affinity to influence the quality and quantity of T cell responses. Indeed, in addition to effects on response magnitude, previous *in vivo* studies have identified divergent differentiation tendencies according to the strength of TCR engagement. While high affinity interactions drive greater accumulation of both effector and memory T cells, effector populations are proportionally larger than those induced by lower affinity interactions; in contrast, weaker TCR stimulation skews T cell phenotypes towards memory precursor, central memory and tissue resident memory states (20,37–43). Accumulating evidence suggests that affinity-dependent differentiation biases promote a high affinity effector response to the current threat, coupled with a broad memory repertoire to defend against mutated pathogen variants (41,44). Several studies have highlighted rapid affinity-dependent phenotypes, such as IRF4 expression and tendency toward asymmetric division, that are associated with altered differentiation outcomes (19,45). Likewise, T cells activated by weaker ligands express less of the autocrine/paracrine growth factor IL2 and mitogenic transcription factor MYC (25,31,46), which promote ribosome biogenesis, proliferation and effector differentiation (47–49). However, key gaps remain in our understanding of how TCR stimulation strength controls differentiation, particularly with respect to early molecular events *in vivo*.

Here, we leveraged an established influenza model of varied TCR-ligand affinity and fate reporter system to comprehensively study the *in vivo* naïve T cell response from initial activation through differentiation. We found that affinity-dependent differentiation state biases affected cells that had followed multiple differentiation routes. Mitogenic activity as opposed to fate markers were the primary axis of affinity-dependent bias early after infection. Computational isolation and comparison of acutely activated, non-dividing cells showed that weakly and strongly stimulated T cells underwent the same TCR-induced transcriptomic changes, but their magnitude was tuned according to ligand affinity. This resulted in high affinity ligands exacerbating TCR-induced suppression of interferon (IFN) response genes and induction of ribosome biosynthesis, while transcription factor binding site enrichment analyses indicated broadly elevated activity of TCR-induced signalling pathways with strong stimulation. Together, these data indicate that the accumulation of TCR-induced signalling products translates TCR binding properties into cell cycle and associated over-representation of effector states *in vivo*.

## RESULTS

### Strong TCR stimulation promotes expansion and short-lived effector fate in lymphoid and non-lymphoid tissues

To alter the strength of TCR stimulation to ovalbumin (OVA) peptide-specific OT-I TCR-transgenic CD8 T cells *in vivo,* we used previously developed influenza A viruses (IAV) that were modified to express variants of the OVA peptide SIINFEKL (5,50). In brief, we adoptively transferred 10^5^ purified naïve OT-I Rag2^−/−^ CD8 T cells into congenic wild-type C57BL/6 hosts before infecting with IAV expressing either high affinity SIINFEKL (N4), ∼5.6-fold reduced affinity SIIQFEKL (Q4) or extremely low affinity EIINFEKL (E1) peptides (51–53) **(Figure 1A**). Analyses of the lung-draining mediastinal lymph node (medLN), spleen and lungs at days 5 and 10 post-infection revealed the greatest abundance of OT-I cells induced by N4-expressing IAV, moderate expansion by Q4-expressing IAV, and minimal expansion by E1-expressing IAV, as previously described (5,50). We noted on day 5 after infection that while N4 induced greater numbers of OT-I cells than Q4 in the medLN, numbers were equal in the lungs and spleen (**Figure 1B**), consistent with a previous report that weakly stimulated T cells undergo more rapid lymph node egress (54). By day 10, cell numbers in the lung and spleen were significantly associated with TCR stimulation strength. Examination of weight loss revealed no consistent differences between mice infected with N4- versus Q4-expressing IAV and weight recovery from day 6 or 7 with all viral variants, indicating that differential OT-I cell abundances were not due to illness severity (**Supplemental Figure 1A**).

**Figure 1.**
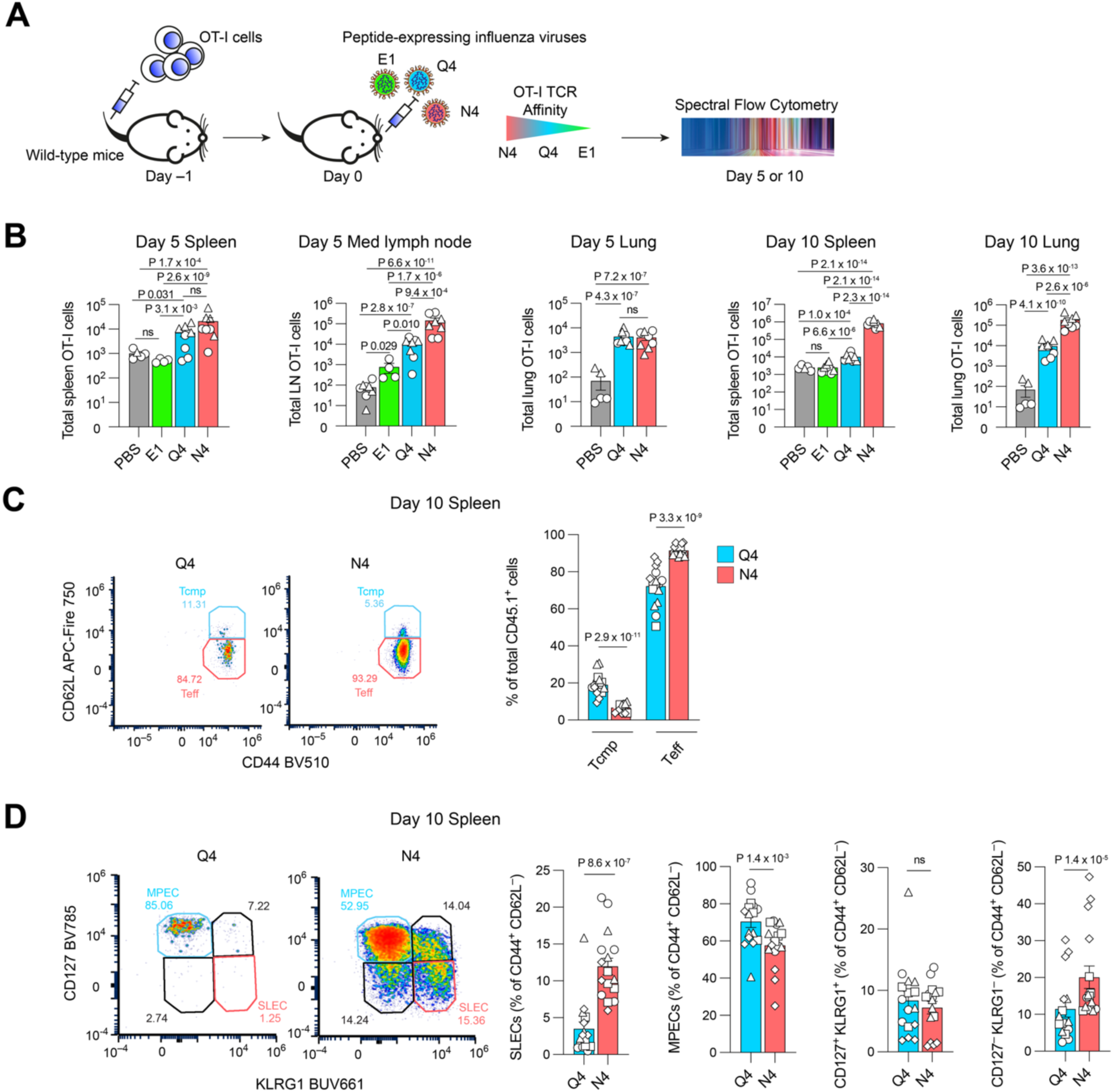
Stimulation strength-dependent CD8 T cell abundance and differentiation fate in the spleen following infection with influenza virus variants. **(A)** Experimental design. CD45.1^+^ OT-I *Rag2*^−/−^ naïve CD8 T cells were adoptively transferred into wild-type CD45.2^+^ hosts before infection with influenza virus variants the next day. Tissues were analysed by flow cytometry on days 5 and 10 post-infection. **(B)** Numbers of adoptively transferred OT-I cells present in various tissues on days 5 and 10 post-infection with influenza viruses; combined data from 2 independent experiments with total 4-8 mice per group. Examining OT-I cells in the spleen at day 10 post infection, frequencies of **(C)** CD62L^high^ T_cmp_ and CD62L^low^ T_eff_ populations and **(D)** among T_eff_, the frequencies of SLEC, MPEC, CD127^+^ KLRG1^+^ and CD127^−^ KLRG1^−^ populations. Numbers on biaxial flow cytometry plots indicate percentages. **(C-D)** Pooled data from 4 independent experiments, each of which included 4 mice per group. **(B-D)** Each point is a mouse; different symbols represent data from independent experiments; bar height and error bars indicate mean ±SEM. As described in Methods, statistical testing accounted for shared donor cells within each independent experiment by including experiment as a covariate in **(B)** 2-way ANOVA without interaction and Tukey’s method for pairwise comparisons and **(C-D)** linear regression.

We next examined the impact of TCR stimulation strength on differentiation state. While all differentiated populations were numerically more prevalent after infection with IAV expressing high (N4) versus reduced (Q4) affinity ligands (**Supplemental Figure 1B-C**), we sought to compare the relative proportions of OT-I cells in each differentiation state according to stimulation strength. We note that any differences in proportional representation of T cell subsets can reflect expansion, cell death, and tissue localisation, as well as differentiation trajectory biases. On day 10 after IAV infection, all splenic OT-I cells expressed elevated CD44, but the proportion of CD62L^high^ cells, indicative of central memory precursors (T_cmp_), was approximately halved after strong (N4) as compared with weaker (Q4) stimulation (**Figure 1C**). Within the CD44^high^CD62L^low^ T_eff_ population, the proportion of short-lived effector cells (SLECs, CD127^−^KLRG1^+^) was increased and the proportion of memory precursor effector cells (MPECs, CD127^+^KLRG1^−^) reduced with strong versus weaker stimulation (**Figure 1D and Supplemental Figure 1D**). Percentages of double-positive effector cells (CD127^+^KLRG1^+^) were equal among N4- and Q4-stimulated OT-I cells, but the percentage of double-negative effector cells, which have capacity to differentiate into either SLECs or MPECs (55) was twice as large after strong N4 stimulation (**Figure 1D**).

As expected for a peripheral tissue, all OT-I cells in the lung were CD62L^low^ and were therefore defined in our gating as T_eff_. Following infection, a population of tissue resident memory T cells (T_rm_) forms in peripheral tissues and can be identified by co-expression of CD103 and CD69. Ten days post-infection, this phenotype likely denotes T_rm_ precursors (T_rmp_). While the total number of T_rmp_ on day 10 post-infection was elevated with strong N4 stimulation (**Supplemental Figure 1C**), the percentage of T_rmp_ among lung OT-I cells was lower after high affinity N4- as compared with reduced affinity Q4-expressing IAV (**Figure 2A**). Gating lung OT-I cells on KLRG1 and CD127 to examine SLEC and MPEC phenotypes showed that, similar to splenic populations, strong stimulation drove proportionally more cells with SLEC and double-negative effector cell phenotypes, and proportionally fewer cells with an MPEC phenotype, compared to weaker stimulation (**Figure 2B**). Examination of the T_rmp_ phenotype within CD127^+^KLRG1^−^ MPECs showed that T_rmp_ constituted on average 44% of total MPECs after weak stimulation but only 20% after strong stimulation (**Figure 2C**). In sum, examination of differentiation states 10 days post infection demonstrated that while strong stimulation increases the number of cells in all examined differentiation states, it also drives in proportionally more cells of a (short-lived) effector phenotype both systemically and at the sight of infection. These data are concordant with results from other *in vivo* models of altered TCR-antigen affinity (20,37–43) and validate use of this influenza model for investigating their upstream molecular and cellular underpinnings.

**Figure 2.**
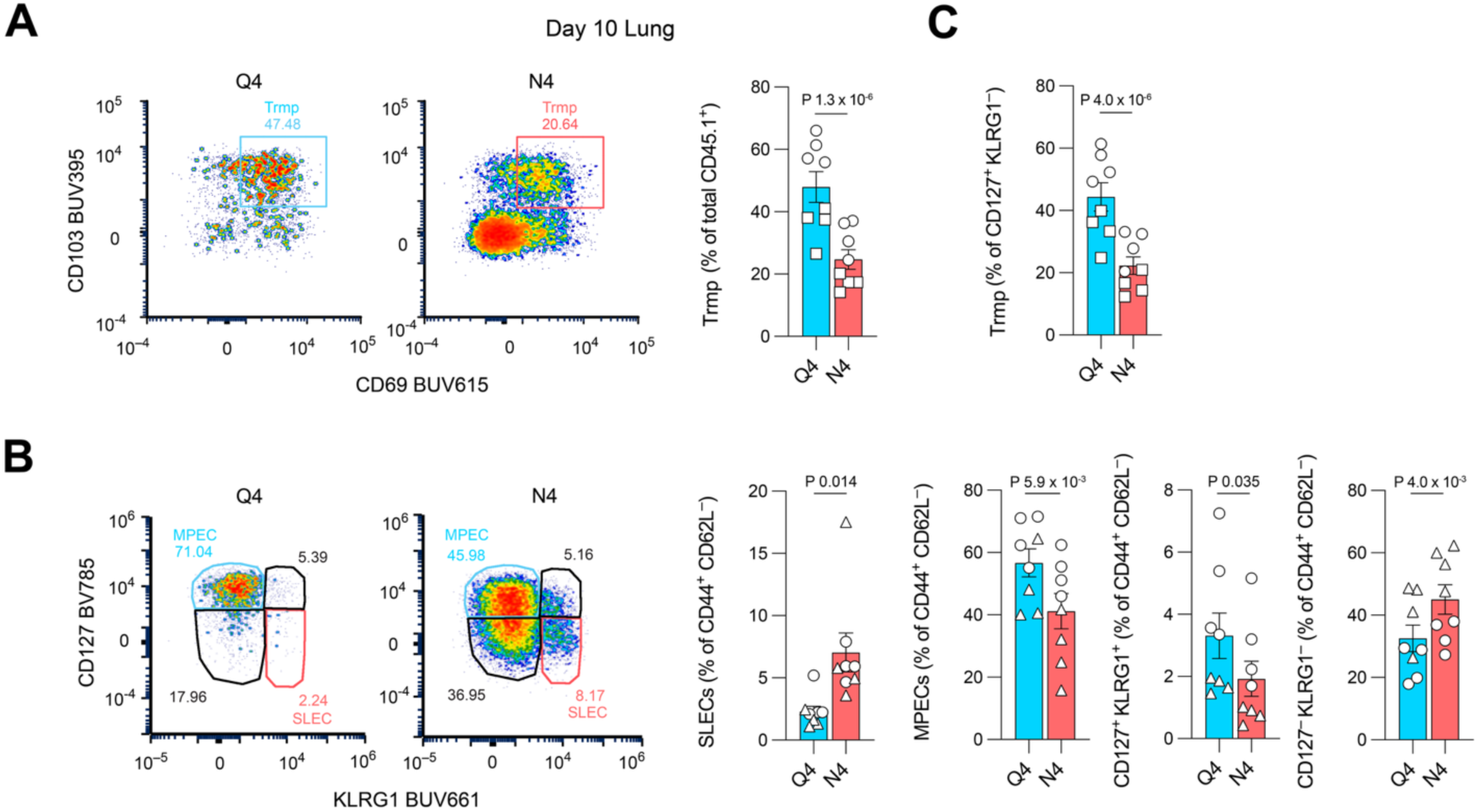
Stimulation strength-dependent differentiation of CD8 T cells in the lung after infection with influenza virus variants. Experimental design as in Figure 1A. Lungs were analysed by flow cytometry on day 10 post-infection. (**A**) The frequency of CD103^+^ CD69^+^ T_rmp_ among OT-I cells. (**B**) Frequencies of SLEC, MPEC, CD127^+^ KLRG1^+^ and CD127^−^ KLRG1^−^populations among OT-I cells which were all of T_eff_ (CD44^high^ CD62L^low^) phenotype. (**C**) Proportion of cells expressing CD103 and CD69 among MPECs. (**A-C**) Combined data from 2 independent experiments, each of which included 4 mice per group. Each point is a mouse; different symbols represent data from independent experiments; bar height and error bars indicate mean ±SEM. As described in Methods, statistical testing was performed by linear regression, accounting for shared donor cells within each independent experiment.

### The impact of TCR-ligand affinity persists through multiple stages of differentiation and protein expression

After initial activation, T cells can embark directly on a memory differentiation trajectory or first adopt an effector phenotype before diverting to a memory fate (14,15). To test whether TCR stimulation strength impacts this process of memory diversion, we used a KLRG1 fate-mapping model, in which cells that express KLRG1, a marker for SLEC phenotype, undergo cre-mediated removal of a transcriptional stop sequence, resulting in constitutive TdTomato expression for themselves and their progeny (15). We transferred naïve OT-I *Rag2*^−/−^*Klrg1*^Cre/+^*Rosa26*^tdTomato/+^ CD8^+^ T cells into congenic hosts before infecting with N4- or Q4-expressing IAV and analysing spleen and lung tissue on day 10 (**Figure 3A**). Similar to previous work with this model, we observed incomplete penetrance of the fate reporter such that only 22-78% of KLRG1^+^ cells expressed TdTomato in the spleen and lung (**Figure S2A**). We also noted that CD44^+^CD62L^high^ T_cmp_ cells were never TdTomato^+^ (**Supplemental Figure 2B**), and thus that diversion of KLRG1^+^ cells to a memory differentiation pathway was limited to the CD62L^low^ MPEC population (CD62L^low^CD44^high^CD127^+^KLRG1^−^) in our model at this time point.

**Figure 3.**
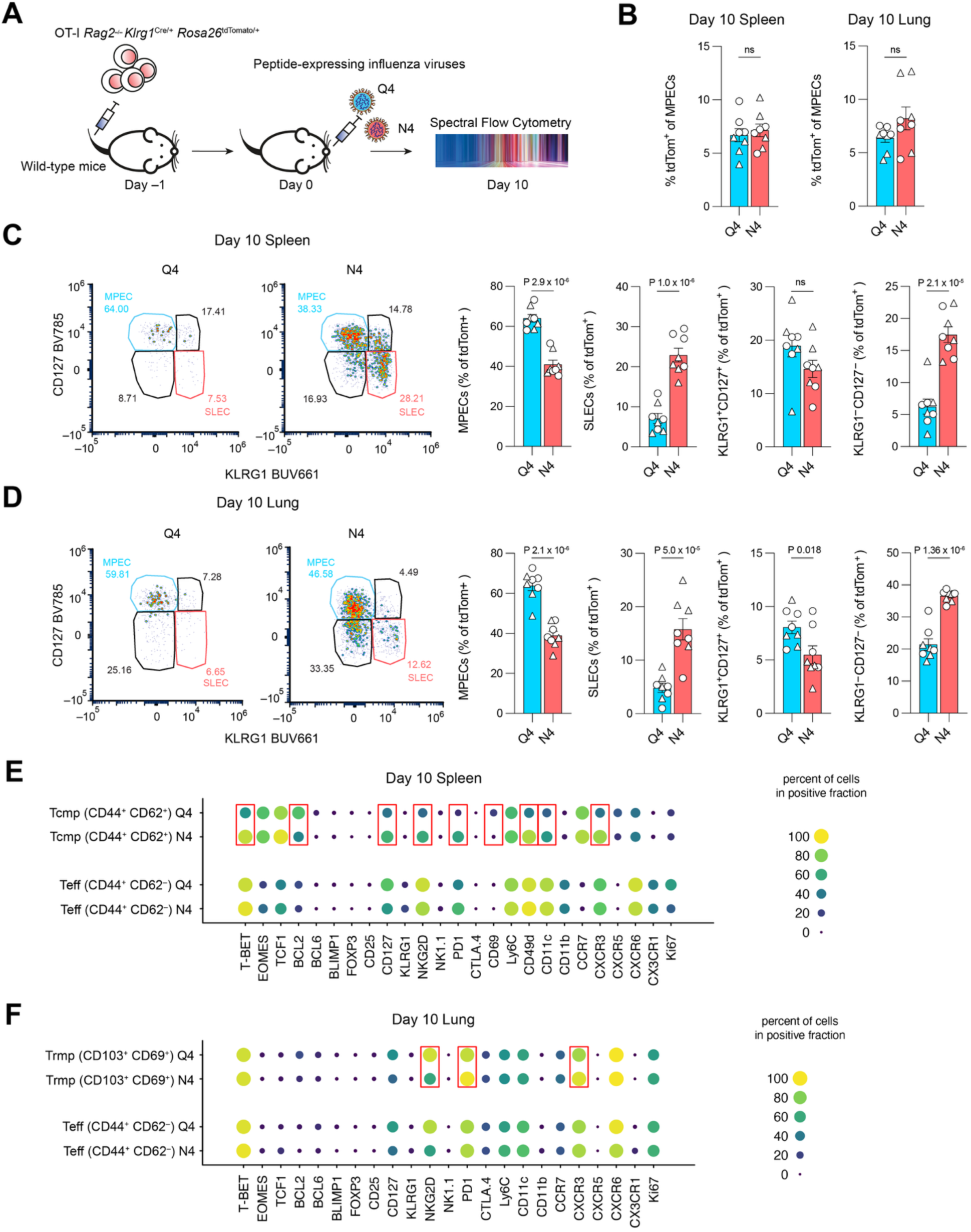
KLRG1 fate-mapping and phenotype comparison among differentiated CD8 T cells after different strengths of TCR stimulation. (**A**) Experimental design to examine memory diversion during differentiation. CD45.1^+^ *Klrg1*^Cre/+^ *Rosa26td*^Tomato/+^ OT-I *Rag2*^−/−^ naïve CD8 T cells were adoptively transferred into wild-type CD45.2^+^ hosts before infection with influenza virus variants the next day. Tissues were analysed by flow cytometry on day 10 post-infection. (**B**) Percentages of tdTomato^+^ (KLRG1 fate-mapped) cells among MPECs in the spleen and lung. Phenotypic analysis of tdTomato^+^ cells in the (**C**) spleen and (**D**) lung. (**A-D**) Combined data from two independent experiments, each of which included four mice per group. Each point is a mouse; different symbols represent data from independent experiments; bar height and error bars indicate mean ±SEM. (**E-F**) Experimental design as in Figure 1A, examining frequency of indicated markers 10 days after infection within differentiated subsets in (**E**) spleen and (**F**) lung; plots depict further analysis of experiments shown in Figures 1C-D and 2, respectively. (**B-F**) As described in Methods, statistical testing was performed by linear regression, accounting for shared donor cells within each independent experiment. (**E-F**) Red box indicates p < 0.05 after Bonferroni correction for the number of markers tested.

Comparing responses to Q4- versus N4-expressing IAVs, we found that 4-12% of current MPECs were TdTomato^+^, regardless of antigen variant, indicating that TCR stimulation strength does not impact the proportion of MPECs derived from each differentiation pathway (**Figure 3B**). To instead examine the fate of all lineages that had passed through a KLRG1^+^ state, we considered the differentiation states of all TdTomato^+^ cells (**Supplemental Figure 2B and C**). This revealed that in both the lung and spleen, TdTomato^+^ cells from strong (N4) stimulation were less likely than those from weaker (Q4) stimulation to be ex-KLRG1 MPECs, instead remaining as KLRG1^+^ SLECs or becoming ex-KLRG1 double negative effector cells (**Figure 3C and D**). These results demonstrate that the proportional skew of weakly stimulated T cells toward memory precursor states also affects cells derived from KLRG1^+^ lineages, driving a persistent bias across cells taking multiple differentiation routes.

We next sought to examine the uniformity of T cells within major defined subsets and test whether expression of key proteins differed according to initial TCR stimulation strength. Leveraging flow cytometry data from Figures 1 and 2 to assess expression of 23-25 additional proteins in T_eff_ and T_cmp_ in the spleen or T_rmp_ in the lung on day 10 post-infection, we identified substantial affinity-dependent protein expression differences within T_cmp_ and T_rmp_ populations (**Figure 3E and F**). For example, on average 50% of weakly (Q4)-stimulated splenic T_cmp_ expressed T-bet, compared with nearly all those receiving strong (N4) stimulation, whereas the opposite was true for the apoptosis regulator BCL2 (**Figure 3E**). These data build upon a previous observation that T-bet induction correlates with ligand affinity and can inhibit transcription of EOMES-induced BCL2 (41) by showing that the expression pattern reflects not only altered differentiation state biases but variation within the T_cmp_ population. T_cmp_ that received a strong stimulus also expressed more NKG2D and PD-1, associated with stronger activation and optimal CD8^+^ T cell memory formation (56,57), as well as cell adhesion and migration-associated proteins CD49d, CD11c and CXCR3, suggesting the potential for differential cell-cell interactions. In the lung, T_rmp_ also showed differential expression of NKG2D, PD-1 and CXCR3 between weakly and strongly stimulated cells, although effect sizes were smaller than in splenic T_cmp_. In marked contrast, T_eff_ had no significant stimulation strength-dependent variability in the proteins examined. Together, these data show that antigen affinity determines not only the percentages of cells that adopt T_cmp_ and T_rmp_ characteristics but also the protein expression phenotypes of these cells.

### Protein expression profiles reflect affinity-associated differentiation biases before the appearance of classical effector subsets

To better understand the origins of differentiation fate biases and the impact of stimulation strength on the early stages of the CD8^+^ T cell response, we examined OT-I cell responses on day 5 after infection. Similar to day 10, the proportion of CD62L^high^ cells was lower after strong (N4) stimulation in the medLN and spleen (**Figure 4A, B**). In the lung, a smaller percentage of strongly as compared to weakly stimulated cells expressed CD69 (**Figure 4C**), which may be an early indicator of their reduced tendency toward tissue residency or reflect different activation kinetics between strongly and weakly stimulated T cells. KLRG1 expression was undetectable at day 5, precluding further delineation of conventional subsets at this time point

**Figure 4.**
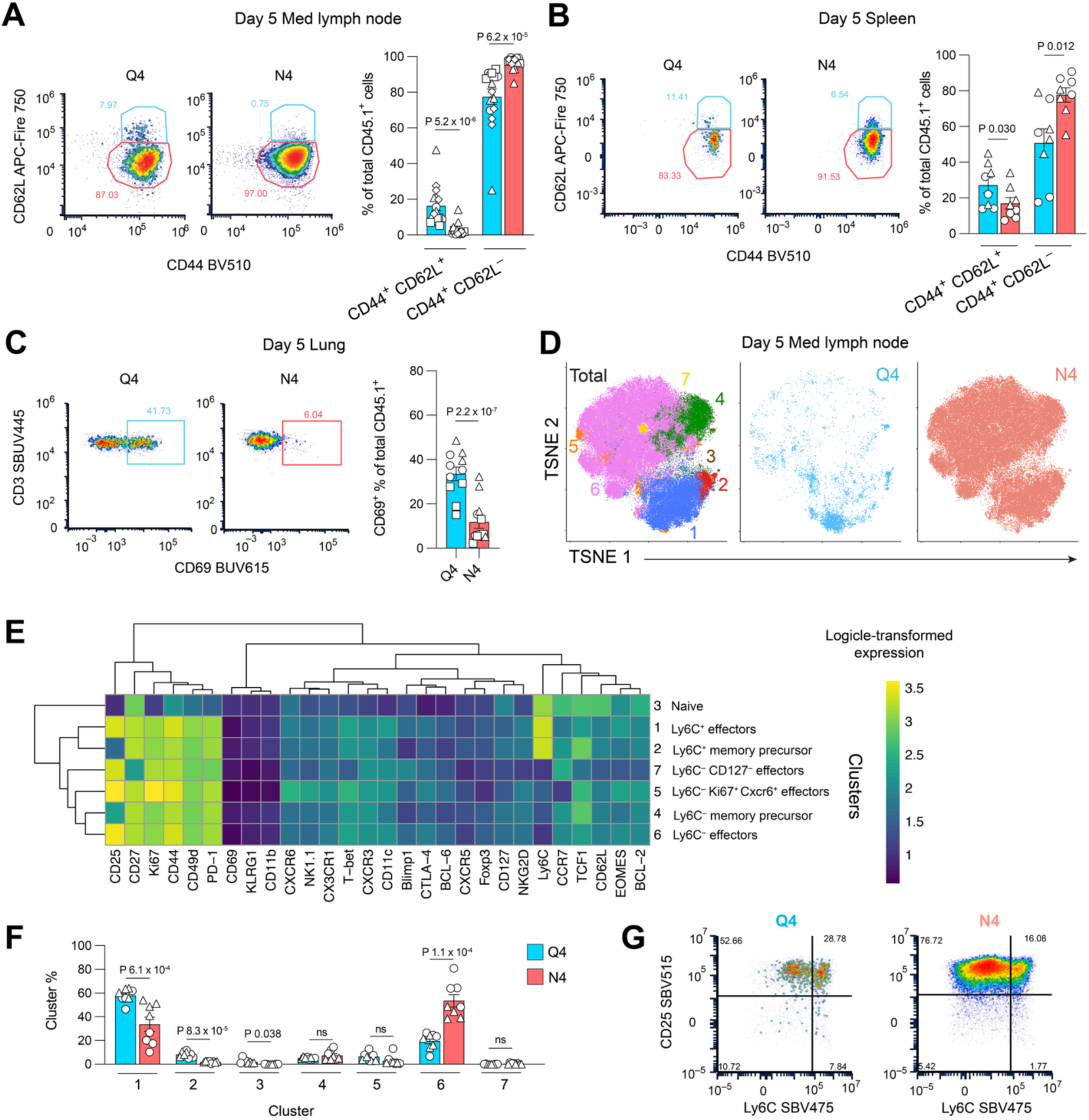
Stimulation strength-dependent phenotypes of CD8 T cells 5 days after infection with influenza viruses. Experimental design as in Figure 1A. Tissues were analysed by flow cytometry on day 5 post-infection. The frequency of CD62L^high^ and CD62L^low^ phenotypes among OT-I cells in (**A**) the medLN and (**B**) the spleen. (**C**) The frequency of CD69^+^ cells among OT-I cells in the lung. (**A-C**) Combined data from 2-4 independent experiments, each of which included 4 mice per group. (**D**) t-SNE representation of OT-I cells from the medLN merged across samples, coloured by cluster (left) and whether they came from mice infected with Q4-expressing IAV (middle) or N4-expressing IAV (right). (**E**) Heatmap of median expression of each protein within each cluster. (**F**) Percentages of cells within each cluster split by viral variant. Data in (**D-F**) are from 1 representative of 2 independent experiments, each of which included 4 mice per group; for (**F**), cells from the second experiment were mapped onto the clusters of the first and plot depicts combined data. (**G**) Representative biaxial plots of Ly6C versus CD25 expression on OT-I cells from the medLN for simplified depiction of data in (**D-F**). (**A-C, F**) Each point is a mouse; different symbols represent data from independent experiments; bar height and error bars indicate mean ±SEM. As described in Methods, statistical testing was performed by linear regression, accounting for shared donor cells within each independent experiment. Bonferroni correction was applied when testing differential abundances across multiple clusters in (**F**).

To characterise early T cell responses in the absence of classical gating makers, we used a self-organising map implemented in FlowSOM (58) to examine the high-dimensional phenotypes of OT-I cells from the medLN at day 5 post infection (**Figure 4D**). Fine-grained clusters were grouped into seven metaclusters, named according to prominent marker expression: naïve cells (c3), two subsets of memory precursors (Ly6C^+^, c2; and Ly6C^−^, c4), two major CD25^+^ effector populations (Ly6C^+^, c1; and Ly6C^−^, c6) and 2 minor effector populations that, in addition to being CD25^+^Ly6C^−^were distinguished by being Ki67^++^CXCR6^+^ (c5) or CD27^−^ (c7) (**Figure 4E**). While not a commonly used T cell marker, Ly6C was previously shown to be expressed at a higher level in naïve CD8 T cells and also inducible by multiple cytokines, particularly type I IFN (59). Thus, the stark Ly6C^+/−^bifurcation we find between otherwise similar populations of activated T cells may indicate altered cytokine signalling activity.

Testing for differential abundances of cell subsets between mice infected with N4- and Q4-expressing IAV revealed greater proportions of naïve (c3), Ly6C^+^ memory precursors (c2), and Ly6C^+^ effectors (c1) with weaker (Q4) stimulation (**Figure 4F**), while strong (N4) stimulation gave rise to more Ly6C^−^ effectors (c6). We were intrigued by the strong inverse association of stimulation strength with Ly6C expression within CD25^+^ effector populations (c1 and c6) (**Figure 4E-G**). Examination of Ly6C expression in naïve CD45.2^+^ host T cells that shared the immune environment with OT-I cells showed no difference after infection with Q4- versus N4-expressing IAV (**Supplemental Figure 2E)**, indicating that differences in OT-I Ly6C expression were not driven by changes in the draining lymph node cytokine milieu but rather were TCR stimulation dependent. Thus, these data show that TCR-ligand affinity-associated biases in differentiation states are detectable on day 5 after infection and that affinity controls expression of Ly6C among activated CD25^+^ cells, suggesting the possibility of differential cytokine response signalling among early effector cells.

### Single-cell RNA-sequencing demonstrates early stimulation strength-dependent biases in coupled cell cycle and metabolic profiles

Based on phenotypic divergence observed by flow cytometry on day 5 post-infection, we next sought to molecularly profile responding T cells as early as possible. To test the feasibility of recovering activated transferred cells from the draining medLN on days 2-5 after IAV infection, we adoptively transferred naïve GFP-*Myc* OT-I cells (which express a GFP-MYC fusion protein from the endogenous *Myc* locus) into congenic hosts before infecting with N4- or Q4-expressing IAV. Recovery of at least a few hundred OT-I cells from the medLN became feasible from day 4 onward, (**Supplemental Figure 2F**), and on days 4 and 5 post-infection we detected GFP^+^ OT-I cells, indicating their TCR activation (**Supplemental Figure 2G**).

We therefore sorted CD45.1^+^ OT-I cells from the medLN of mice 4 and 5 days after infection for single-cell RNA-sequencing (scRNA-seq). Oligonucleotide hashtags allowed pooling of cells from mice infected with both N4- and Q4-expressing viruses, thereby avoiding confounding of stimulation strength with technical batch. The transcriptomes of cells from each day and virus infection were heterogeneous but showed clear condition-specific enrichment of phenotypes when visualised in 2-dimensional t-SNE plots (**Figure 5A**). We used graph-based clustering to group cells according to transcriptome similarity and annotated these clusters using cluster-associated marker genes (**Supplemental Table 1**) and key genes from the literature, identifying naïve/weak activation, acute TCR stimulation (characterised by the highest expression of TCR-induced genes including *Myc,* and *Nr4a1*), central memory precursors, two clusters of non-cycling early effector cells, and 5 clusters of cycling early effector cells (**Supplemental Figure 3A**).

**Figure 5.**
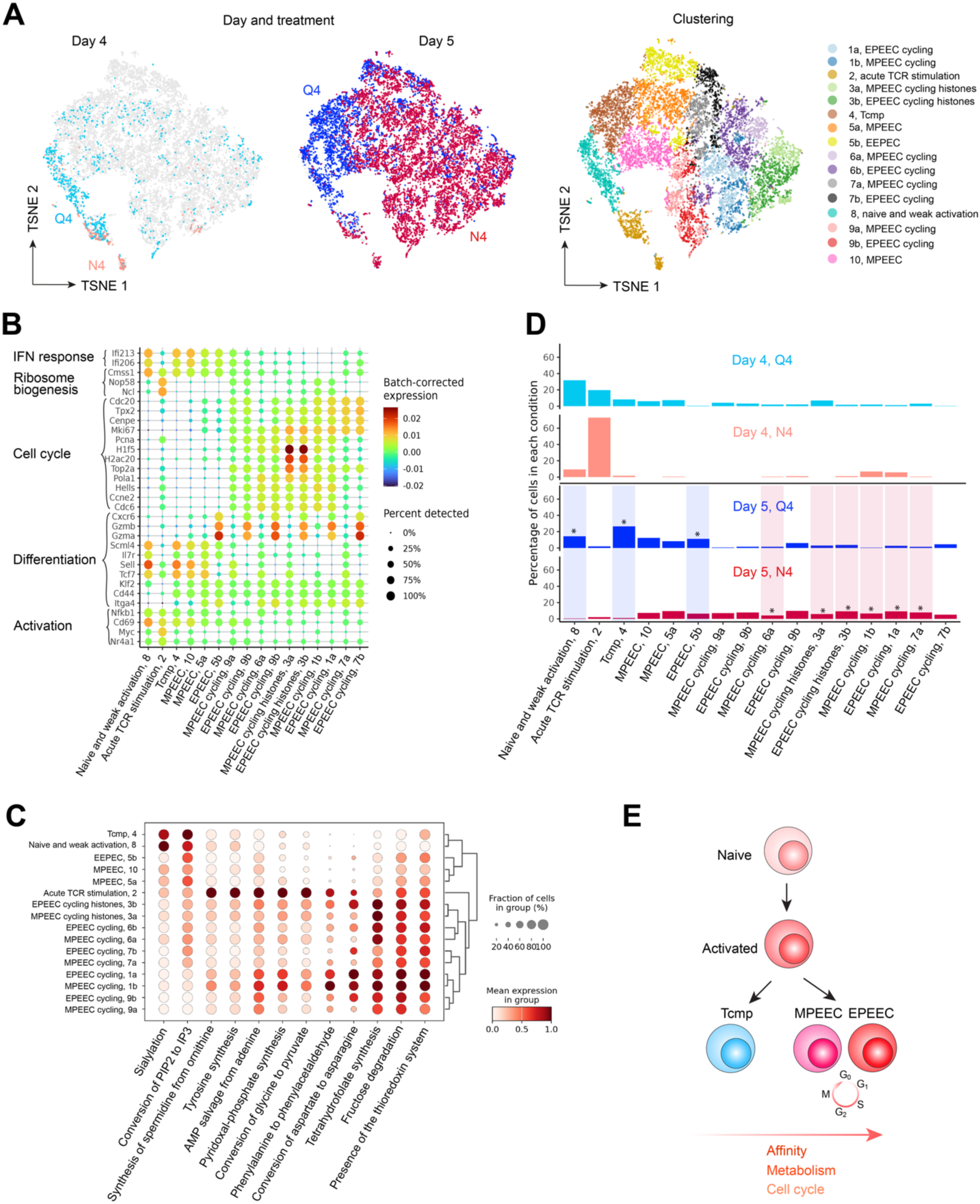
scRNA-seq analysis of CD8 T cells following infection with influenza viruses expressing peptide antigens with high or reduced TCR affinity. CD45.1^+^ GFP-MYC OT-I *Rag2*^−/−^ naïve CD8 T cells were adoptively transferred into wild-type CD45.2^+^ hosts before infection with Q4- or N4-expressing influenza viruses the next day. OT-I cells were sorted from medLNs on days 4 and 5 post-infection. (**A**) t-SNE plots coloured by viral variant on day 4 (left) and day 5 (middle) post-infection and transcriptomic cluster (right). (**B**) Dot plot illustrating expression of variably expressed genes used for cluster annotation. Dot colours depict batch-corrected, normalised log-expression. (**C**) Dot plot illustrating metabolic tasks that showed the statistically significant differences in inferred activity between clusters (see Methods). (**D**) Cluster occupancy of cells from each viral variant and day. Differential abundance testing was performed between viruses separately on days 4 and 5. Statistically significant differences (FDR < 0.05) are indicated by an asterisk over and shading in the colour of the population that is significantly greater. (**E**) Cartoon summarising differentiation pathways and associations with antigen affinity and cell cycle and metabolic activity from our data.

The marker genes for many clusters were dominated by cell cycle-associated genes; while this is an important feature of early T cell activation (60), high variance in transcripts related to cell cycle can mask early markers of differentiation fates. We therefore examined *Tcf7* (encoding TCF1) as a key indicator of memory potential and performed subclustering on those clusters with bimodal *Tcf7* expression. This split all cycling cell states into memory-phenotype early effector cell (MPEEC) and effector-phenotype early effector cell (EPEEC) subsets (**Figure 5B**), characterised by divergent expression of fate-associated genes such as *Tcf7, Sell, Gzma* and *Gzmb* (**Supplemental Figure 3B**).

Some of the hallmarks of T cell activation and differentiation are shifts in key metabolic pathways that facilitate biosynthetic processes and rapid proliferation (61,62). Newly developed tools for scRNA-seq data analysis, such as scCellfie (63), use prior knowledge of reactions, enzymes, and their encoding genes from genome-scale metabolic models to infer the activity of metabolic tasks from single-cell transcriptomic data, facilitating deeper inferences into cell state. As would be expected, scCellfie analysis among the T cell clusters showed evidence of greater metabolic activity across a broader range of pathways in cycling early effector cells as compared with central memory precursor or naïve T cells (**Supplemental Figure 4A, Supplemental Table 2**). Examination of significantly differentially active tasks between clusters (see Methods) revealed a particularly striking elevation in “Tetrahydrofolate synthesis” in cycling early effector subsets (**Figure 5C**), consistent with the importance of one-carbon metabolism in the synthesis of nucleic acids and cell division (64). Interestingly, MPEEC and EPEEC subsets of the same early effector clusters showed near-identical metabolic task activities (**Supplemental Figure 4B**). Statistical comparison of metabolic task activities in MPEEC versus EPEEC subsets within each cluster revealed very few tasks specifically associated with EPEEC or MPEEC subsets, with minimal overlap between clusters (**Supplemental Figure 4C**), indicating that there is no metabolic phenotype that separates cycling early effector cells that express genes associated with effector versus memory fates. These data indicate that metabolic state is strongly associated with cell cycle status, but that early effector cells with the same cell cycle and metabolic profiles can vary in their expression of fate-associated genes.

Cells within the “acute TCR stimulation” cluster showed the highest level of activity in the largest number of differentially active metabolic tasks, specifically including the strongest evidence for upregulation of “synthesis of spermidine from ornithine”, “tyrosine synthesis”, “AMP salvage from adenine”, “pyridoxal-phosphate synthesis” (PLP synthesis), and “conversion of glycine to pyruvate” (**Figure 5C**). Several of these processes reflect amino acid metabolism pathways, which have been previously identified as key components of TCR-induced T cell activation (65–67), while the AMP salvage pathway provides a rapid source of nucleotides often in the context of rapidly growing and expanding cells (57). The unique nature of the active metabolic tasks in this cluster suggests a short window during T cell activation when specific biosynthetic switches are required to progress activated T cells into the next stage of proliferation and differentiation.

Finally, we tested for differential abundances of cells within each cluster between N4- and Q4-expressing virus infections, separately on days 4 and 5 (**Figure 5D, Supplemental Table 3**). We first noted that cells from each IAV variant were present in each cluster, demonstrating that all states were possible with either stimulus. No significant biases in subset abundances were observed on day 4, likely influenced by high variability in the number of cells recovered per mouse at this time point. On day 5, Q4-stimulated OT-I cells were strongly skewed toward the naïve/weak activation or T_cmp_ clusters, with additional subtle enrichment in non-cycling EPEECs. In contrast, N4-stimulated cells were more likely to be in cycling early effector clusters, notably within both EPEEC and MPEEC subsets. These data indicate that fate biases directed by TCR stimulation strength manifest at day 5 in the medLN as unequal distributions of metabolically distinct subsets: cycling early effectors associated with strong TCR stimulation, and non-cycling central memory precursor and early effector cells associated with activation by weaker TCR stimulation.

### TCR-induced transcript abundance is quantitatively tuned by ligand affinity *in vivo*

We postulated that comparing the transcriptomes of cells showing signs of recent TCR stimulation after N4- versus Q4-expressing IAV infection could reveal the early consequences of affinity-dependent differential signalling. Because of the inherent response time heterogeneity among T cells during *in vivo* infection, we reasoned that it was more reliable to identify recently activated cells by transcriptomic state than by post infection time point. We therefore used a pseudobulk differential expression analysis to compare strongly (N4-) versus weakly (Q4-) stimulated cells within the “acute TCR stimulation” cluster of our scRNA-seq data from days 4 and 5 post infection (**Figure 6A, Supplemental Table 4**). Among the significantly differentially expressed genes, we found many that were previously reported to reflect the strength of TCR stimulation, such as *Il2* (25,69), *Irf4* (32,70,71), *Xcl1, Ccl3* and *Ccl4* (34), associated with strong N4 stimulation, and *Tapbp* and *Ifi47* (34), associated with weaker Q4 stimulation (**Figure 6A,B**). Gene ontology enrichment analyses revealed that strongly stimulated T cells exhibited higher expression of genes involved in cell division T cells, while weakly stimulated cells preferentially expressed genes associated with T cell activation and signalling (**Figure 6C, Supplemental Table 5**).

**Figure 6.**
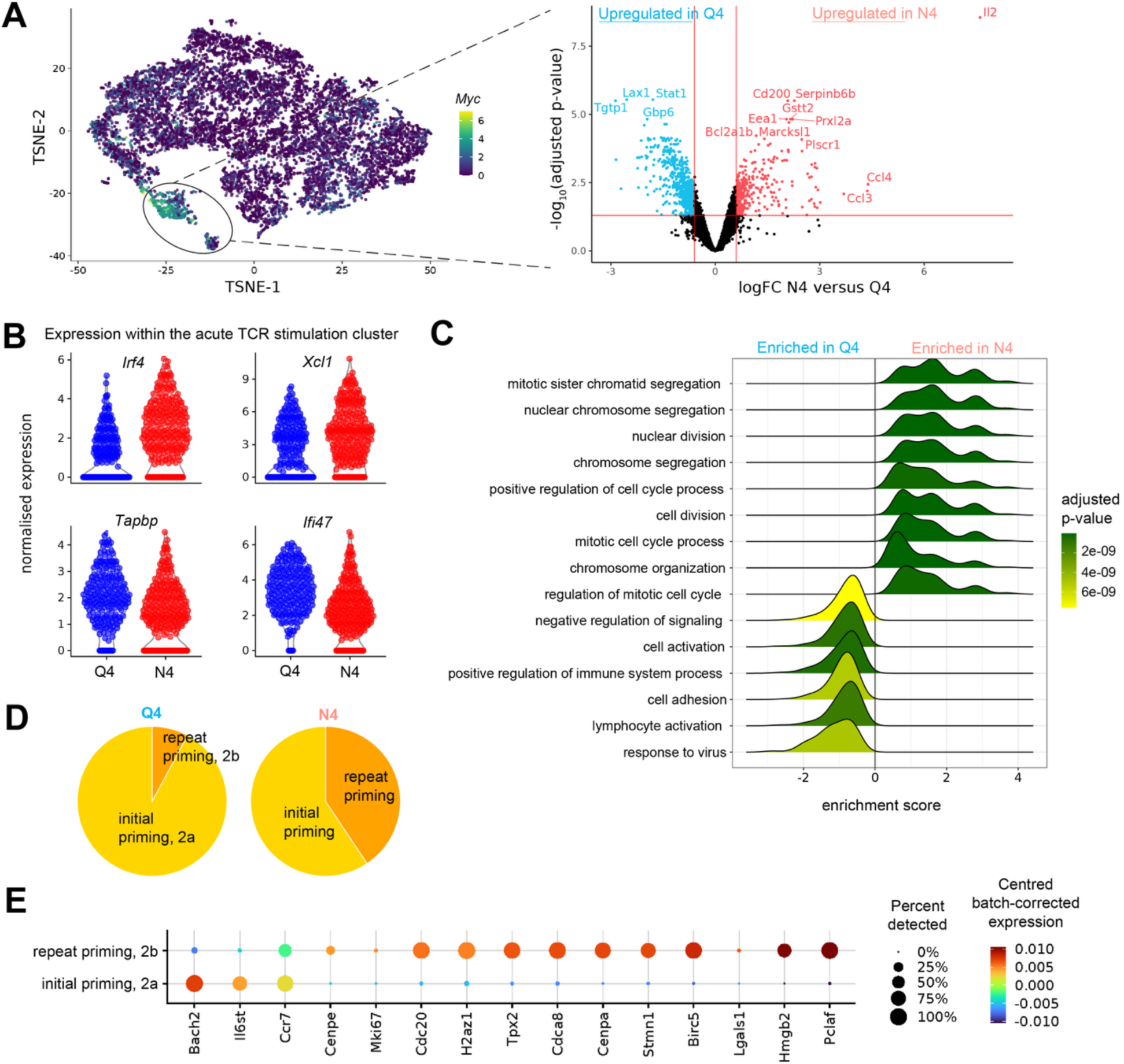
Transcriptome comparison of CD8 T cells acutely stimulated with high versus reduced affinity ligands during influenza infection. (**A**) T-SNE of scRNA-seq data from Figure 5 (OT-I cells sorted from the medLN on days 4 and 5 post-infection with N4- and Q4-expressing IAV) coloured by *Myc* expression and highlighting cluster 2, “acute TCR stimulation” (left). Volcano plot depicts results of a pseudobulk differential expression analysis between cells from N4- versus Q4-expressing IAV infections in the “acute TCR stimulation” cluster (right); positive log-fold-change (logFC) indicates higher expression with N4 stimulation. (**B**) Selected significantly (FDR < 0.05) differentially expressed genes; y-axis is normalised log-counts. (**C**) Ridge plot shows GO category enrichment of differentially expressed genes; positive enrichment score indicates enrichment with N4-expressing IAV. (**D**) Pie charts showing the proportion of cells in 2a, “initial priming”, and 2b, “repeated priming”, subclusters from Q4- and N4-expressing IAV infection conditions. (**E**) Dot plot of variably expressed genes used to annotate subclusters “initial” and “repeated priming” subclusters.

Naïve T cells are quiescent and only initiate cell cycle progression after approximately 24 hours from TCR stimulation. Therefore, concurrent expression of genes indicative of acute TCR stimulation with cell cycle genes suggests repeated priming, as recently described by Jobin *et al.* (36). To limit our investigation to T cells most likely to be in their first round of priming, we subclustered the “acute TCR stimulation” cluster into non-cycling cells, more likely to be in an “initial priming” phase, and cycling cells, likely to be undergoing “repeated priming”. Notably, the “repeated priming” phenotype was more prevalent after high affinity N4 stimulation, concordant Jobin *et al.*’s observation that strongly stimulated T cells undergo a second priming phase (36) (**Fig 6D,E**).

To characterise the earliest transcriptomic effects of altered TCR stimulation strength *in vivo*, we first used pseudobulk differential expression analyses to examine how strongly and weakly stimulated “initial priming” cells differed from those of a “naïve and weak activation” phenotype. Comparison of effect sizes revealed strong correlation, as would be expected because the N4-and Q4-stimulated “initial priming” cells had been clustered together (**Figure 7A, Supplemental Table 4**). However, fitting a trendline to this effect size comparison demonstrated a striking deviation from equality: strong N4 stimulation drove on average 1.3 times more extreme transcriptome remodelling than weaker Q4 stimulation along the shared response axis (**Figure 7A**). This trend of affinity-dependent exaggeration of TCR-induced changes in transcript abundance affected nearly all genes, both up- and down-regulated (**Figure 7A**), with the notable exception of *Il2,* which was selectively expressed by cells undergoing “initial priming” with N4 stimulation (**Figure 7B**). This pro-proliferative and pro-effector cytokine was previously shown to be dispensable for early TCR-induced cell cycling *in vivo* but important for prolonged expansion (72,73) and may therefore create delayed autocrine feedback to drive proliferation and effector fate. Taken together, these data indicate that early after *in vivo* activation, the vast majority of TCR-induced transcriptomic changes are quantitatively controlled by the strength of TCR engagement.

**Figure 7.**
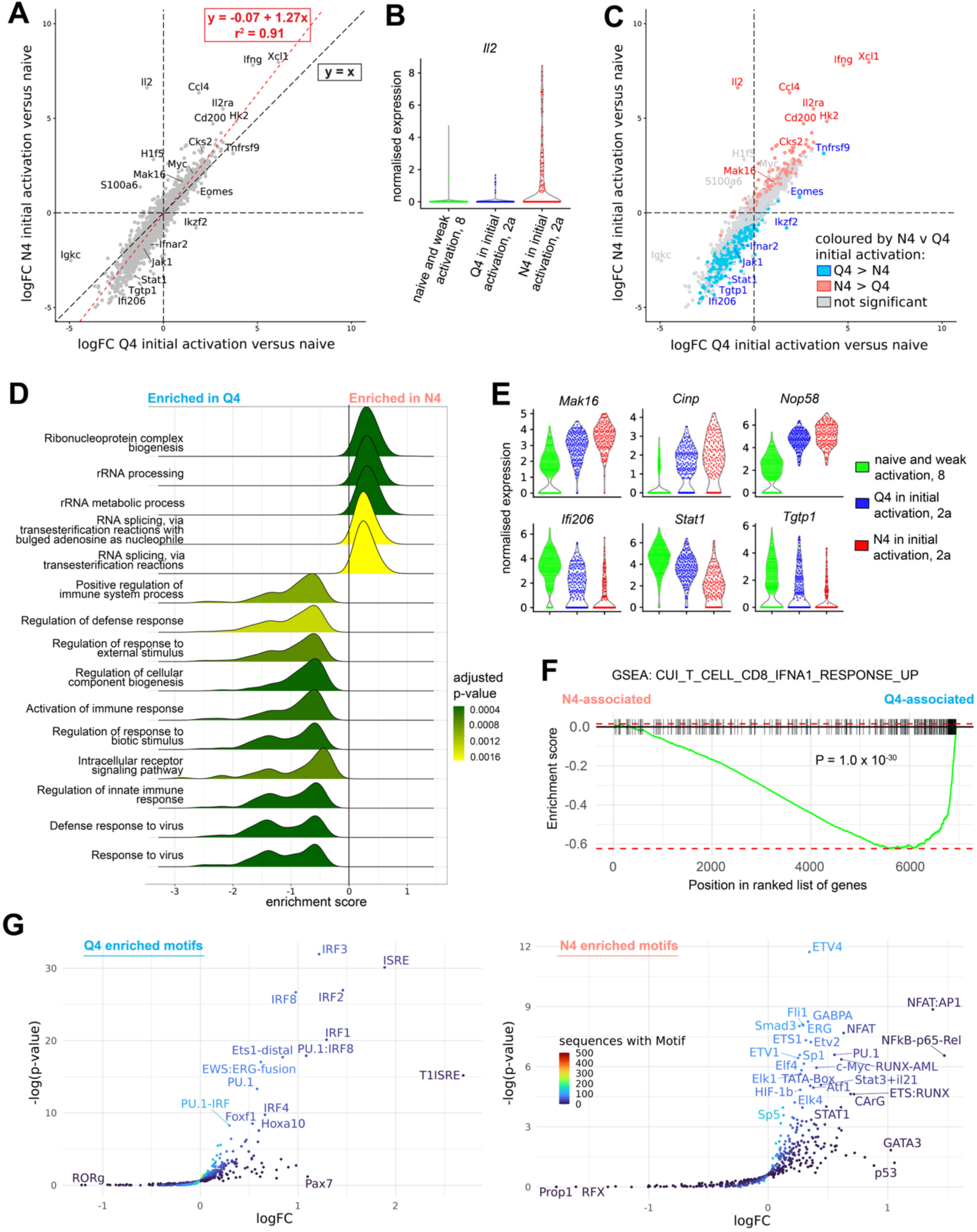
Transcriptome and signalling inference in CD8 T cells undergoing initial priming *in vivo* with strong versus reduced affinity ligands. Analyses of cells from the non-cycling “initial priming” subcluster of the “acute TCR stimulation” cluster. (**A**) Log-fold-change plots comparing results of differential expression analyses between N4-stimulated or Q4-stimulated cells from the “initial priming” subcluster versus all cells in the “naïve and weak activation” cluster. Red line and equation from linear model fit to the data. (**B**) *Il2* expression in the “naïve and weak activation” cluster, and the “initial priming” subcluster split according to Q4- or N4 stimulation; y-axis is normalised log-counts. (**C**) Plot as in (**A**) coloured by results of direct differential expression analysis between N4- and Q4-stimulated cells from the “initial priming” subcluster. (**D**) Ridge plot shows GO category enrichment of differentially expressed genes between N4 or Q4 “initial priming”; positive enrichment score indicates enrichment with N4-expressing IAV. (**E**) Expression of selected ribosome biogenesis (top) or interferon pathway (bottom) genes in the “naïve and weak activation” cluster and the “initial priming” subcluster split according to Q4- or N4 stimulation; y-axis is normalised log-counts. (**F**) Gene set enrichment analysis of differentially expressed genes between N4 or Q4 “initial priming” as in (**D**) using a bespoke set composed for genes upregulated in CD8 T cells in response to IFNα stimulation. (**G**) Volcano plots depict enrichment of transcription factor binding sites among differentially expressed genes between N4- and Q4-stimulated cells from the “initial priming” subcluster.

### Strong TCR stimulation exaggerates inverse regulation of ribosome biosynthesis and anti-viral response genes

Direct differential expression and gene ontology enrichment analyses between N4- and Q4-stimulated cells undergoing “initial priming” highlighted the cellular processes most impacted by quantitative transcriptome remodelling: ribosome biogenesis was the most upregulated and viral response the most downregulated by strong stimulation (**Figure 7C-E, Supplemental Table 4**). Ribosome biogenesis is required for TCR-induced proliferation and is regulated by the coordinated activity of TCR, costimulatory and IL2 signalling, mediated at least in part by MTORC1 (25,74). To confirm via an orthogonal measurement that TCR-ligand affinity affects this pathway, we stimulated naïve OT-I cells with N4 or Q4 peptides *in vitro* for 1 hour before measuring phosphorylation of the S6 ribosomal subunit. In accordance with previous results, we found that pS6 levels were dependent on ligand affinity in acutely activated cells *in vitro* (**Supplemental Figure 5A**) (25,33).

In contrast to induced ribosome biogenesis, TCR ligation suppressed viral response genes in a manner dependent on ligand affinity. Many anti-viral genes are components of the type I IFN signalling pathway, which is known to have anti-proliferative activity and be dampened by TCR engagement (75,76), but has not been explored in the context of altered stimulation strength. To specifically investigate this pathway, we ran an additional gene set enrichment analysis using a curated list of genes induced by type I IFN stimulation in CD8 T cells and revealed significantly higher expression with weaker Q4 as compared to stronger N4 initial priming (**Figure 7F**). These data are concordant with our day-5 flow cytometry observation that Ly6C expression remained higher in activated cells after infection with Q4-expressing versus N4-expressing IAV (**Figure 4E-H**). Moreover, T cells stimulated *in vitro* in the presence of exogenous IFNα confirmed that Ly6C expression is suppressed in a manner dependent on TCR-ligand affinity (**Supplemental Figure 5B**). Together, these data highlight the coordinated, inverse regulation of ribosome biogenesis and IFN response gene expression, with TCR stimulation strength dictating the size of this trade-off.

Finally, to understand the origins of these gene expression differences in T cells recently activated by viruses expressing strong (N4) versus weaker (Q4) ligands, we looked for enrichment of transcription factor (TF) binding sites among the significantly differentially expressed genes (**Figure 7G, Supplemental Table 6**). The top “initial priming” Q4-enriched TF binding sites included IRF3, ISRE and IRF2, concordant with evidence of greater IFN signalling. In contrast, “initial priming” N4-enriched TF binding sites comprised multiple TCR-induced TFs, including ETS family members, NFAT:AP1, NFAT and NFkB-p65-Rel, which are activated downstream of MAP kinases (ERK, JNK, p38), calcium and NF-κB pathways (3,77). These data indicate that strong TCR stimulation amplifies the gene expression outputs of the TCR-induced signalling network by quantitatively enhancing TF activity. A corollary of this conclusion is that cells stimulated by N4 and Q4 ligands employ the same pathways to achieve a biosynthetic pro-effector state and that Q4 ligation is simply, on average, less efficient in achieving this. Accordingly, *in vitro* T cell activation in the presence of the clinically approved allosteric MEK1/2 inhibitor Trametinib confirmed that ERK1/2 signalling is required for induction of effector-associated proteins and cell cycle initiation (78–80), and showed that this is independent of TCR stimulation strength (**Supplemental Figure 5C-G**). Taken together, our examination of T cell priming *in vivo* demonstrates that strong TCR ligation broadly amplifies TCR-induced signalling network activity and gene expression outputs to efficiently promote a coordinated pro-mitogenic state driving expansion and propensity toward effector states (**Supplemental Figure 6**).

## DISCUSSION

Our molecular and phenotypic investigation of varying TCR stimulation strength *in vivo* adds several new layers to our understanding of how TCR-ligand interactions are translated into functional outcomes. We found that TCR-ligand affinity impacts not only numbers and proportions of cells in effector and memory precursor differentiation states but also fate distributions of cells from KLRG1+ lineages and phenotypes of differentiating memory precursor populations. Early after activation, we uncovered a divergence in cell cycle and metabolic properties dependent on TCR stimulation strength and showed that this can be explained by quantitative tuning of TCR-induced transcriptome remodelling.

While previous studies showed that TCR stimulation strength can bias differentiation fate distributions (20,37,38,41–43), it has remained unclear whether this represents a simple probabilistic mechanism or whether phenotypes of cells within defined subpopulations are also affected. Our data indicate the latter among splenic T_cmp_ and lung T_rmp_ at day 10 after infection. Such regulation of differentiated cell properties beyond subset membership is concordant with a previous report of differential transcript expression between T_rm_ derived from high or reduced affinity TCR stimulation in IAV infection (38). These observations of affinity-associated phenotypes in memory and memory precursor populations but not in effector cells raises the intriguing possibility that effector tuning may be primarily quantitative while memory tuning is also qualitative.

Using a KLRG1 fate mapping model, we demonstrated that the memory precursor state bias of weakly stimulated cells exists both in initial commitment to effector versus memory precursor fate and in later diversion from SLEC to MPEC trajectories. This may reflect affinity-dependent differences in expansion of SLECs that maintain KLRG1 expression or in the probability of embarking on the memory diversion differentiation trajectory. Considering the latter possibility in conjunction with the recent observation from Jobin *et al.* that strongly stimulated cells can undergo additional priming raises the possibility of a second round of TCR-driven differentiation decisions. Moreover, strongly stimulated T cells are known to spend more time in the secondary lymphoid tissue (23,54), potentially facilitating additional rounds of TCR stimulation that could amplify fate biases as they proliferate.

An intriguing observation from our scRNA-seq data is that while T_cmp_ clearly separate from early effector cells based on fate markers, cell cycle and metabolic profiles, there is also a bifurcation of early effector cells with memory versus effector phenotypes (MPEEC versus EPEEC) in all stages of the cell cycle. Asymmetric division is a tempting explanation, but evidence for this mechanism has mainly been reported in contexts where metabolic mediators also divide unequally (49,81), which we do not find in MPEEC versus EPEEC comparisons in our data. Instead, our results highlight that biosynthetically active, cycling early effector cells can exhibit signs of memory fate, dissociating fate markers from mitogenic properties.

Through differential expression analyses in cells showing signs of initial priming *in vivo*, we found that high affinity ligand engagement led to global exacerbation of TCR-induced transcriptome remodelling. These data build upon previous findings that TCR stimulation strength tunes the magnitude of TCR-induced changes in transcript abundance *in vitro* (29,30) to show that that this occurs in a complex *in vivo* setting. At the extreme end of affinity-dependent downregulation of transcript expression, we found IFN pathway genes. It remains an intriguing focus for future studies to better understand why TCR signalling suppresses IFN response genes and how this relates to the antiproliferative effects of IFN (75,76,82). In contrast, at the extreme end of affinity-dependent upregulation, we found ribosome biogenesis, which is critical for activated T cells to enter the cell cycle (25). Rapid cycling is closely associated with effector differentiation fates, with some studies suggesting a causal role (60,83,84), and thus early pro-mitogenic activity is concordant with the subsequent prevalence of cycling early effectors and short-lived effector differentiation states among strongly stimulated cells.

TF binding site enrichment analysis among transcripts most highly expressed with high as compared with reduced affinity priming identified canonical TCR-induced pathways including ERK/MAPK, NFAT and NF-κB. While many TCR-induced signalling events exhibit digital (all-or-nothing) behaviour after ligation, a recent *in vitro* study using live-cell signalling reporters over time showed more nuanced dynamics such that ERK1/2 and NFAT signalling persisted and/or repeatedly pulsed with strong but waned with weaker TCR stimulation (85). Moreover, finely controlled experiments using an optogenetic chimeric antigen receptor system showed that abundances of induced transcripts reflected the persistence of receptor signalling in T cells (80). TCR-induced transcript levels could thereby act as a time-integrated measure of TCR engagement, which is longer and/or more frequent with higher affinity ligands. Our data tie these concepts to the functional T cell response *in vivo* by demonstrating that the degree of transcriptome remodelling downstream of TCR signalling foreshadows affinity-dependent expansion and differentiation state biases.

*Il2* transcripts were found almost exclusively among T cells receiving a strong stimulus during initial priming. How this is achieved at the signalling level remains a key open question. IL2 coordinates with TCR signalling to promote ribosome biogenesis and biosynthetic activity via MTORC1 and MYC in activated cells (25,31,74). Indeed, supplementation with exogenous or paracrine IL2 disproportionately enhances proliferation in weakly stimulated cells (25,46,86). While previous work showed that autocrine IL2 is dispensable for early rounds of division after *in vivo* TCR stimulation (72,73), its affinity-dependent expression suggests a route for subsequent amplification of affinity-driven biases in biosynthetic activity and associated effector fate (87–89).

A limitation of our study is the use of a TCR-transgenic system: this allowed us to tightly control TCR stimulation strength in an *in vivo* respiratory infection, but future development of systems in which it is possible to reliably measure both antigen affinity and molecular properties in individual cells will be important to determine generalisability of these findings.

In conclusion, our investigation of how TCR-ligand affinity impacts CD8 T cell differentiation *in vivo* showed that high affinity TCR-ligand interactions amplify TCR-induced transcriptomic changes driving IFN response downregulation and cell cycling, followed by a persistent short-lived effector state bias and altered phenotypes among memory precursor cells. Our results suggest that quantitative gene expression tuning allows TCR-induced signalling machinery to record the duration and/or frequency of TCR-ligand interactions in each naïve T cell, controlling initiation of mitogenic processes that ultimately drive expansion and associated differentiation states. Such a mechanism would enable a high degree of tuning according to antigen binding properties and persistence, facilitating host responses of appropriate antigen specificity, magnitude and function.

## MATERIALS AND METHODS

### Study Design

The objective of our study is to investigate how antigens of different TCR affinities influence differentiation decisions taken by CD8^+^ T cells following activation. Purified naïve CD8^+^ T cells from CD45.1 Rag2^−/−^ OT-I mice (8 - 24 weeks-old) were adoptively transferred into sex and age-matched CD45.2 C57/BL6 recipient mice (8 - 12 weeks-old). Mice were randomly assigned to either the control or experimental groups prior to the start of the experiment to eliminate potential litter and cage biases. All experimental groups had a minimum of four animals per group. The day after adoptive transfer, recipient mice were intranasally challenged with influenza viruses expressing variants of ovalbumin peptide of known affinity for the OT-I transgenic TCR. The number of adoptively transferred OT-I T cells was optimized to ensure adequate recovery of transferred cells even under the conditions of limited expansion due to weak antigen stimulation. Transferred cells were analysed in and recovered from the lung, spleen and/or medLN. Phenotypic changes were measured at two different time points (days 5 and 10) by spectral flow cytometry. To explore transcriptional changes occurring early following *in vivo* activation, cells recovered from medLNs were subjected to scRNA-seq on days 4 and 5 post infection. No data points were excluded, and all experiments were performed on at least two independent occasions, indicated in the figure legends. Investigators were not blinded to the collected data as measurements taken were objective.

### Animals

Wild-type C57BL/6, OT-I *Rag2*^−/−^ (Tg(TcraTcrb)^1100Mjb^ *Rag2^tm1Fwa/tm1Fwa^*), GFP-*Myc* OT-I *Rag2*−/−*(Tg(TcraTcrb)*^1100Mjb^ *Rag2^tm1Fwa/tm1Fwa^ Myc^tm1Slek^) and OT-I Klrg1^Cre/+^ Rosa26^tdTomato/+^ (Tg(TcraTcrb)*^1100Mjb^ *Rag2^tm1Fwa/tm1Fwa^ KLRG1^Cre/+^ Gt(ROSA)26Sor^tm14(CAG-tdTomato)Hze/+^)* mice were bred in-house at the Babraham Institute’s animal facility under specific pathogen-free conditions. All animal experiments were approved by the ethical committee of the Babraham Institute, and conducted according to local guidelines and UK Home Office regulations under the Animals Scientific Procedures Act 1986 (ASPA), Home Office License PP9973990. All mice were fed standard mouse chow ad libitum and were maintained in individually ventilated cages with a 12-hour light/dark cycle.

### Adoptive transfer

For adoptive transfer, naïve CD8^+^ OT-I T cells were purified from the spleen and lymph nodes of OT-I Rag2^−/−^ mice by negative selection using the EasySep Mouse Naïve CD8^+^ T cell Isolation Kit according to the manufacturer’s instructions (Stem Cell Technologies, cat #19858). Purified cells were resuspended in RPMI media supplemented with 1% foetal calf serum. Purity (>90%) was confirmed by flow cytometry before intravenous injection. Each mouse received 10^5^ purified CD8^+^ OT-I T cells.

### Influenza infection

Influenza A viruses (HKx31) that express SIINFEKL (N4), SIIQFEKL (Q4) or EIINFEKL (E1) peptides, described previously (5) were propagated in embryonated chicken eggs. Viruses were diluted in PBS prior to intranasal administration. Mice were anesthetized and infected with 10^2^ PFU.

### Antibodies

Flow cytometric staining was performed by incubating cells with antibodies against CD49d (R1-2), CD115 (T38-320), CD69 (H1.2F3), KLRG1 (2F1), CD27 (LG.3A10), CD11b (M1/70), CCR7 (4B12), pS6 (N7-548) from BD Biosciences; CD3 (KT3), Ly6C (ER-MP20), CD25 (PC61.5.3), CD8 (KT15) from Bio-Rad; CD45.2 (104), NKG2D (CX5), EOMES (Dan11mag), Foxp3 (FJK-16s), Ki67 (SolA15) from Thermo. Fisher; BCL-6 (REA373), T-bet (REA102) from Miltenyi; TCF1 (C63D9), S6 (54D2), p44/42 MAPK (Erk1/2)(Thr202/Tyr204) (D13.14.4E) from Cell Signaling; CD45.1 (A20), CXCR5 (L138D7), CD44 (IM7), CD4 (GK1.5), CTLA-4 (UC10-4B9), CXCR3 (CXCR3-173), CXCR6 (SA051D1), CD11c (N418), CD127 (A7R34), MHCII (M5/114.15.2), CD19 (6D5), PD1 (29F.1A12), CD64 (X54-5/7.1), NK1.1 (S17016D), BCL-2 (BCL/10C4), Ly6G (1A8), Blimp (5E7), CD62L (MEL-14), CX3CR1 (SA011F11), CD3 (145-2C11), CD8 (53-6.7) from Biolegend. Hashtagging for 10X Genomics was performed using TotalSeqB anti-mouse Hashtag 1-6 (M1/42; 30-F11) antibodies from Biolegend.

### Sample preparation for flow cytometry

To obtain single-cell suspensions, finely-chopped lung tissue was digested with 720 mg/ml Collagenase D (Roche) in RPMI media at 37 °C for 30 minutes. The digest was then passed through a 70 μm cell strainer before being centrifuged through 30% Percoll Plus (GE Healthcare, cat # GE17-5445-01) in RPMI. Spleens and lymph nodes were disrupted by mushing through 70 μm cell strainers. Spleens and lungs were subsequently subjected to red blood cell lysis solution (eBioscience, cat # 00-4300-54). Single-cell suspensions from all tissues were resuspended in PBS / 2% FCS. Cells were counted using a Countess cell counter (Thermo Fisher Scientific).

### Flow cytometry

Approximately 2 million cells from each sample were stained for flow cytometry. Nonspecific binding was blocked using 2.4G2 hybridoma supernatant, after which cells were stained with surface antibodies and fixable viability dye ViaKrome 808 (Beckman Coulter, cat #c36628) for 1 hour at 4 °C. Cells were either analysed immediately or fixed and permeabilized using the eBioscience Foxp3 / Transcription Factor Staining Buffer Set (Thermo Fisher Scientific) according to manufacturer’s instructions. For intracellular staining, cells were incubated with antibodies overnight at 4 °C. Flow cytometry samples were acquired on either a Fortessa (BD Biosciences) or an Aurora (Cytek) spectral flow cytometer. Data was analysed using FlowJo (Tree Star, Inc.) or FCS Express (De novo) software packages.

### High dimensional spectral flow cytometry data analysis

Raw flow cytometry data was pre-processed in FCS Express, cleaning the data using FlowCut, scaling, and extracting CD45.1^+^ (adoptively transferred) cells. Clustering was then performed using in R (v4.4.2) using the flowCore (2.18.0) and FlowSOM (2.14.0) packages. After logicle transformation, the FlowSOM algorithm was used without scaling to create a self-organising map and 7 metaclusters.

### Cell sorting and scRNA-seq

Four and five days post-infection, medLNs were dissected and single cell suspensions were prepared as described above. Cells were stained with CD45.1-PE, CD45.2-APC, CD8-BV711, DAPI and one hashtag antibody per mouse. CD45.1^+^ CD8^+^ T cells were then sorted on a FACSAria (BD Biosciences). For each Chromium 10x capture reaction, sorted cells from 6 mice (3 N4- and 3 Q4-infected) were multiplexed together. Three independent experiments were performed, using the following batch structure: batch 1 = day 4; batch 2 = 1 pool day 4, and 1 pool day 5; batch 3 = 1 pool day 4, and 1 pool day 5.

10x Genomics single cell library prep, relevant quality control and sequencing was carried out by the Babraham Institute Genomics Facility. Cells were processed to Gel Beads in Emulsion (GEMs) using the Chromium iX Controller. Following manufacturers guidelines and input cell stock concentration, the maximum number of cells was targeted per sample. Following the GEM generation the library preparation for the whole transcriptome (gene expression) and hashtag libraries was completed, following manufacturer’s instructions, using the Chromium GEM-X Single Cell 3ʹ Kit v4 (PN-1000692) and 3’ Feature Barcode Kit (PN-1000262). Library QC steps were completed using an Agilent 4150 Tapestation with the High Sensitivity D1000 Assay (Agilent, Santa Clara, USA) for the cDNA generation, cDNA amplification and final libraries. Quantification of the final libraries was determined by qPCR with the QuantaBio sparQ Universal Library Quant Kit (QuantaBio). Libraries were pooled and sequenced on the Element Biosciences AVITI platform using a high output sequencing kit (Illumina).

### Single-cell RNA-seq data analysis

Fastq files were demultiplexed and aligned using Cell Ranger multi (Cell Ranger v9.0.1), with the GRCm39 genome as reference. Subsequent processing used R or python. Genes were filtered for minimum expression within each batch and treatment group whilst cells were filtered for minimal reads, minimal *Malat1* expression and extreme mitochondrial gene expression. We noted that day 4 N4-stimulated cells were the most frequently filtered due to negligible *Malat1* expression, and this was not batch-specific. The dataset was checked for CD8 T cell purity. Batch correction was performed using fastMNN (90) (batchelor v1.20.0). Cells underwent unsupervised clustering using Jaccard distances and Louvain community detection on a Shared Nearest Neighbours graph based on batch corrected coordinates, to generate 10 primary clusters. Clusters 1, 3, 5, 6, 7, and 9, and latterly 2, were then subclustered via the same method using batch corrected highly variable genes specific for each cluster and differing k values.

Within-cluster differential expression analysis between N4- and Q4-stimulated cells was performed using a pseudobulk approach, combining cells from the same cluster and sample. These data were fitted with a negative binomial generalised log-linear model (NB-GLM) including batch as a covariate, and differential expression was assessed using a quasi-likelihood F-test, in edgeR (v4.2.0). Of note, where N4- and Q4-stimulated cells from the same “initial priming” subcluster were compared to the “naïve and weak activation” cluster, only log-fold changes were considered because the pseudobulk samples compared were derived from clusters defined by differential expression and some host mice contributed cells to multiple pseudobulked samples. Differential abundance testing was similarly performed using a NB-GLM followed by quasi-likelihood F-tests in edgeR (with trend estimation turned off). Gene ontology enrichment analysis was performed using clusterProfiler(v4.12.6) to examine differentially expressed genes (false discovery rate (FDR) < 0.05) using the Bioconductor genome wide annotation for mouse to provide GO annotation (org.Mm.eg.db v3.19.1). Bespoke gene set enrichment analysis to examine type I IFN-induced genes was performed with fgsea (v 1.32.4) using the CUI_T_CELL_CD8_IFNA1_RESPONSE_UP dataset from MiSigDB. Transcription factor binding site enrichment analysis among differentially expressed genes was performed using Homer v4.11.1 (91) to look for known transcription factor binding site motifs among the promoters of differentially expressed genes. Homer was set to default parameters (a promoter region of –300 to 50 and the default promoter set for mouse).

To compare metabolic activity between clusters we used scCellFie (v0.4.4) (63). For this analysis, cluster assignments, celltype annotation and all metadata were retained but batch correction was re-performed using the Scanpy (v1.10.3) integration of MNN Correct (90). Metabolic activities were inferred within each cluster, and pairwise differential analyses comparing between clusters were performed using the Wilcoxon rank sum test: for Figure 5C, we plotted all tasks with FDR < 0.05 and logFC ≥ 1.5 in at least one pairwise comparison, after removing redundant terms; for Supplemental Figure 4B, an FDR < 0.05 and Cohen’s D equal to or greater than 0.75 were used to indicate significance.

### Cell culture conditions, stimulation and inhibition assays

Cells were cultured in RPMI media supplemented with 10% foetal bovine serum, 50 μM β-mercaptoethanol, 2 mM L-glutamine, 10 units/mL penicillin/streptomycin and 1mM sodium pyruvate at 37°C in a 5% CO2 incubator. All of the above reagents were from Gibco, Thermo Fisher Scientific. OT-I cells were activated by stimulation with either Q4 or N4 peptide at 10nM (Cambridge Bioscience). Where indicated, IFNα (R & D Systems) was added at 1000 U/ml or Trametinib (Selleck Chemicals) at 0.03 μM. All assays were performed over 24 hours.

### Statistical Analysis

Cell numbers in Figure 1B were log-transformed and fitted with an analysis of variance model including experimental replicate and viral variant as covariates. P-values for comparing between peptides were calculated using a 2-way ANOVA without interaction term and Tukey’s method to account for multiple testing in pairwise comparisons. All other N4 versus Q4 comparisons, were made by fitting a linear model including experimental replicate and viral variant as covariates. Bonferroni correction was used to account for multiple comparisons where indicated in the figure legend.

## Acknowledgments

We thank Steve Turner for providing OVA peptide-expressing influenza viruses, Richard Flavell and Ishigame Harumichi for the KLRG1-Cre mouse strain, and Louise Webb and George Smith for valuable feedback on the study. We also thank the Biological Support Unit, Flow cytometry, Genomics, and Bioinformatics facilities at the Babraham Institute for enabling this work.

## Author contributions

Conceptualization: ACR

Methodology: OB, EA, JQ, DL, SC, MT, RV-T

Investigation: VP, AM

Visualization: VP, AM, EA

Funding acquisition: ACR, MT

Project administration: ACR

Supervision: ACR, AL, RV-T

Writing – original draft: VP, ACR

Writing – review & editing: EA, DL, MT, RV-T, ACR

