## Supplemental Figures for "T cell receptor-ligand affinity quantitatively tunes transcriptome remodelling *in vivo* inversely regulating cell division and interferon response"

### Affiliations:

**Supplemental Figures:**

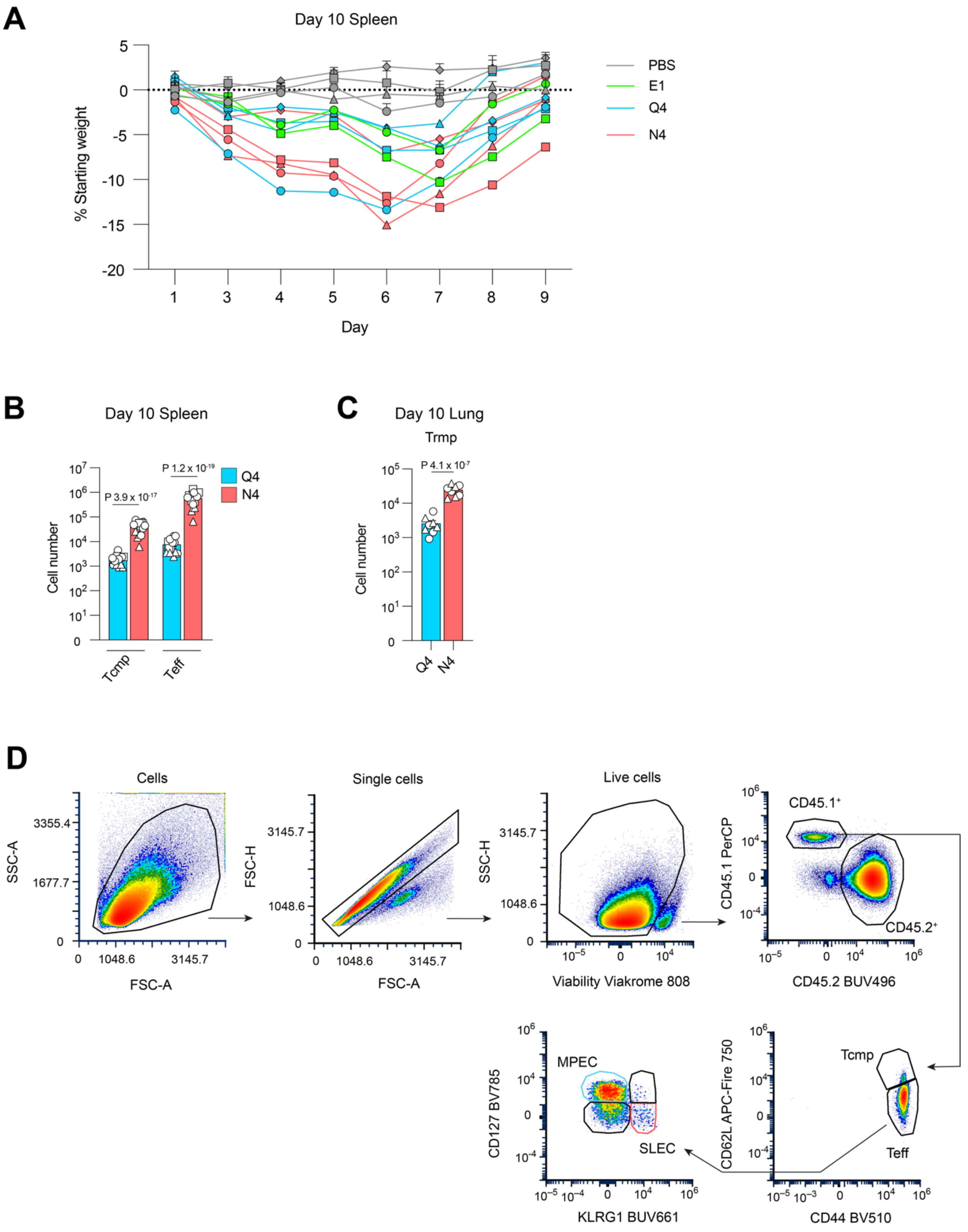

**Supplemental Figure 1. Weight loss during influenza virus infection, OT-I cell numbers in the spleen and the lung, and gating strategy for OT-I T cell sub-populations.** Experimental design as in Figure 1A. (**A)** Weights of host mice that received adoptively transferred OT-I cells followed by infection with either E1-, Q4- or N4-expressing IAV or no infection PBS control. Combined data from 4 independent experiments where each line represents the average across 4 mice from the same condition in each independent experiment. Error bars indicate mean ±SEM. (**B**) Numbers of CD62L$^{high}$ $T_{cmp}$ and CD62L$^{low}$ $T_{eff}$ populations among OT-I cells in the spleen on day 10 post infection. (**C**) Numbers of CD103$^{+}$ CD69$^{+}$ $T_{rmp}$ among OT-I cells in the lung on day 10 post infection. (**B-C**) Combined data from 2-4 independent experiments, each of which included 4 mice per group. Each point is a mouse; different symbols represent data from independent experiments; bar height and error bars indicate mean ±SEM. As described in Methods, statistical testing was performed by linear regression, accounting for shared donor cells within each independent experiment. (**D**) Gating strategy for adoptively transferred OT-I cells and their sub-populations.

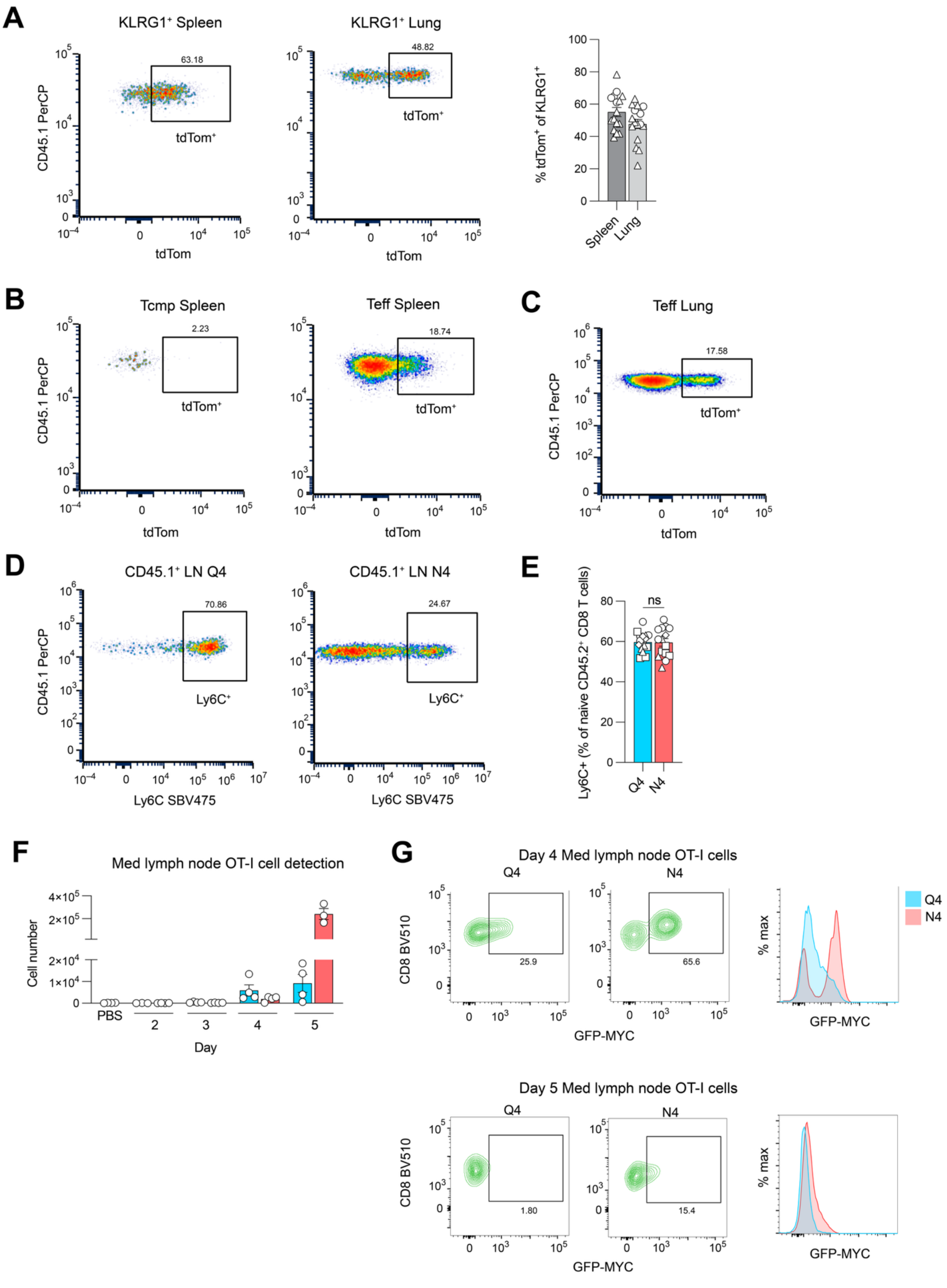

A
KLRG1+ Spleen
KLRG1+ Lung
63.18
48.82
tdTom+
CD45.1 PerCP
tdTom
% tdTom+ of KLRG1+
Spleen
Lung
B
Tcmp Spleen
Teff Spleen
2.23
18.74
C
Teff Lung
17.58
D
CD45.1+ LN Q4
CD45.1+ LN N4
70.86
24.67
Ly6C+
Ly6C SBV475
E
ns
Ly6C+ (% of naive CD45.2+ CD8 T cells)
Q4
N4
F
Med lymph node OT-I cell detection
Cell number
PBS
Day
G
Day 4 Med lymph node OT-I cells
Day 5 Med lymph node OT-I cells
CD8 BV510
GFP-MYC
25.9
65.6
1.80
15.4
% max

**Supplemental Figure 2. Expression of tdTomato in KLRG1 fate-mapping model and analysis of tissues from GFP-MYC mice following infection with influenza viruses expressing high or reduced affinity TCR ligands.** (**A**) Proportion of cells expressing tdTomato of total KLRG1$^+$ cells. Numbers on biaxial flow cytometry plots indicate percentages of pre-gated KLRG1$^+$ population. Graph contains combined data from from two independent experiments, each of which included four mice per group. Different symbols represent data from independent experiments; bar height and error bars indicate mean ±SEM. (**B-C**) Gating for tdTomato$^+$ cells among CD62L$^{high}$ $T_{cmp}$ and CD62L$^{low}$ $T_{eff}$ populations in the spleen (**B**) and CD62L$^{low}$ $T_{eff}$ in the lung (**C**) on day 10 post infection. (**D-E**) Experimental design as in Figure 1A. (**D**) Gating for Ly6C$^+$ cells among CD45.1$^+$OT-I cells in the spleen on day 10 post infection. (**E**) The frequency of Ly6C$^+$ cells among naïve CD45.2$^+$ host CD8 T cells. Combined data from 4 independent experiments, each of which included 4 mice per group. Each point is a mouse; different symbols represent data from independent experiments; bar height and error bars indicate mean ±SEM. (**F**) Absolute numbers of OT-I cells in the mediastinal lymph nodes of mice infected with either Q4- or N4-expressing IAV on days 2, 3, 4 and 5 post-infection as compared to uninfected control mice. Data from one experiment with 3-4 mice per group. (**G**) Expression of GFP-MYC in OT-I cells from (**F**) on day 4 (top) and 5 (bottom) of infection with recombinant influenza viruses.

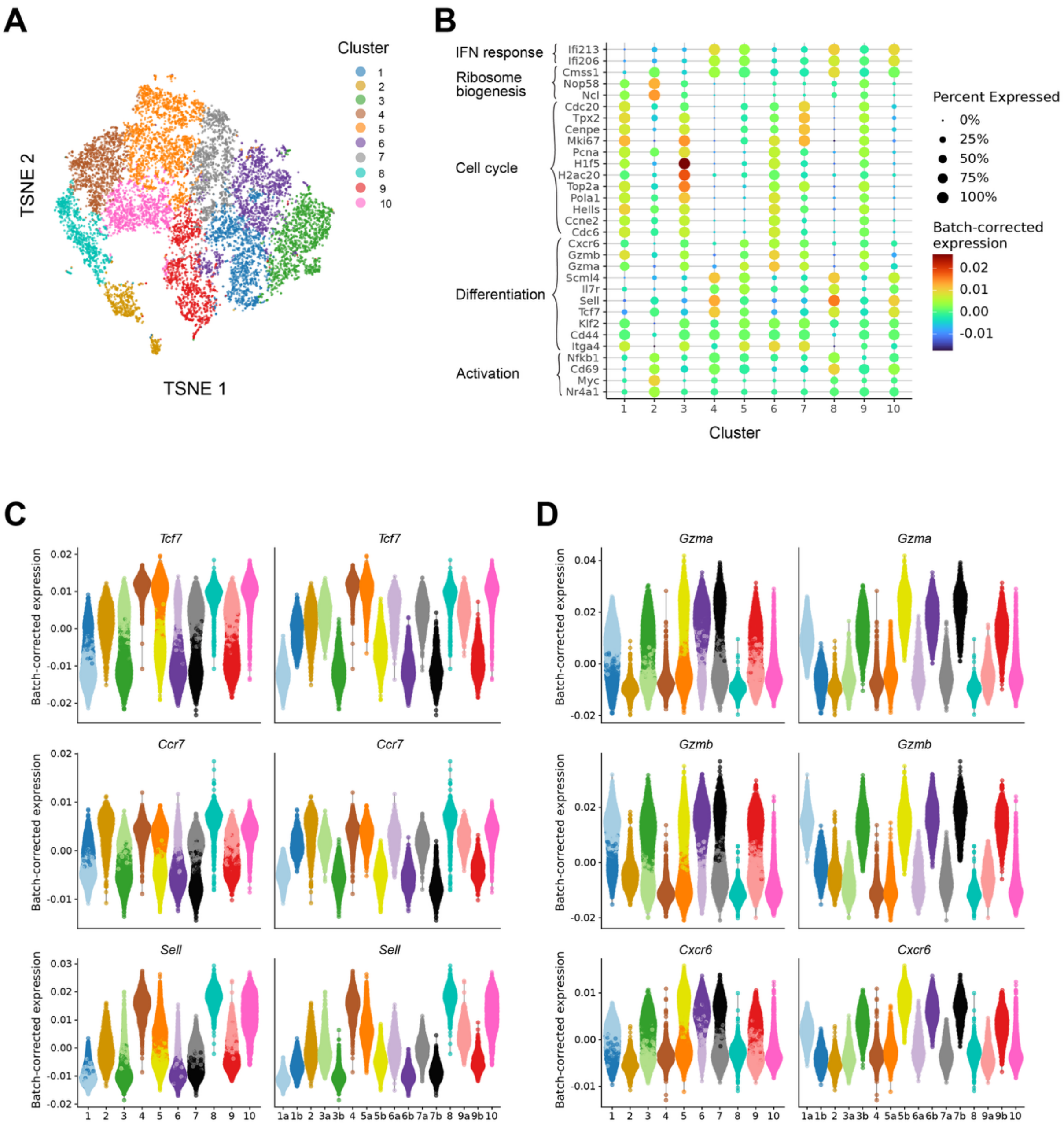


**Supplemental Figure 3. Subclustering of single-cell RNA-seq data from CD8 T cells responding to high or reduced affinity TCR ligands during influenza virus infection.** (**A**) t-SNE plot coloured by initial clustering of OT-I cells. (**B**) Dot plot illustrating expression variably expressed genes used for initial cluster annotation. Dot colours depict batch-corrected, normalised log-expression. (**C-D**) Expression of selected fate-associated genes across (**C**) the initial clusters and (**D**) subsequent subclusters, coloured according to subclusters.

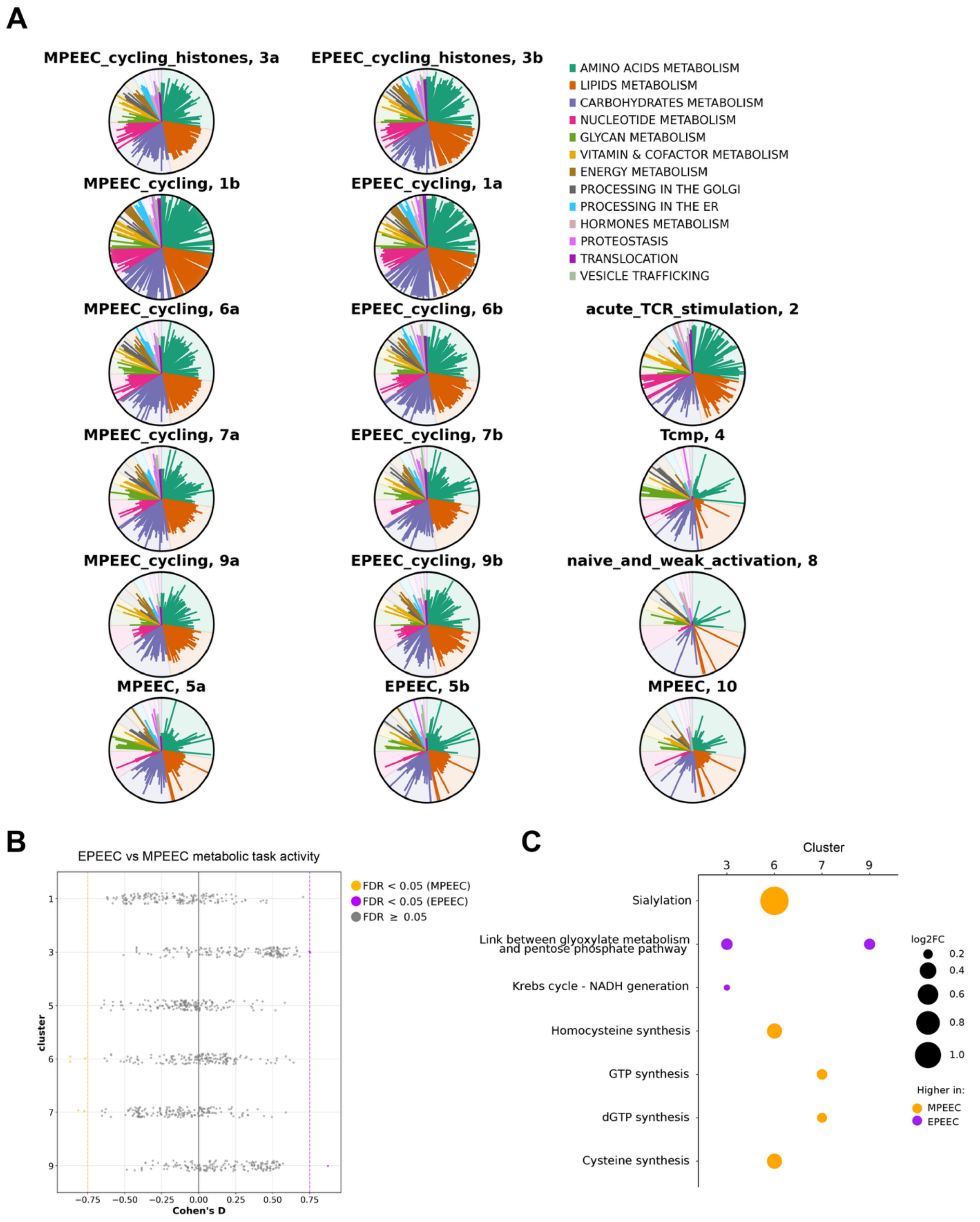


**Supplemental Figure 4. Metabolic task activity in scRNA-seq clusters.** (**A**) Radial plots depict metabolic task activity in each cluster of scRNA-seq data from Figure 5. Each bar represents a metabolic task. Bars originate from the centre, with lengths indicating activity in the indicated cluster relative to the maximum activity detected in any cluster; tasks are colour-coded according to functional grouping. (**B**) Results of differential analyses comparing metabolic task activity in MPEEC versus EPEEC subclusters for each primary cluster containing these cell types (see Methods). Each dot represents a metabolic task, repeated for each y-axis cluster; x-axis shows the effect size of differential activity. (**C**) Dot plot of all metabolic tasks exhibiting statistically significant differential activity in MPEEC versus EPEEC subclusters as highlighted in (B).

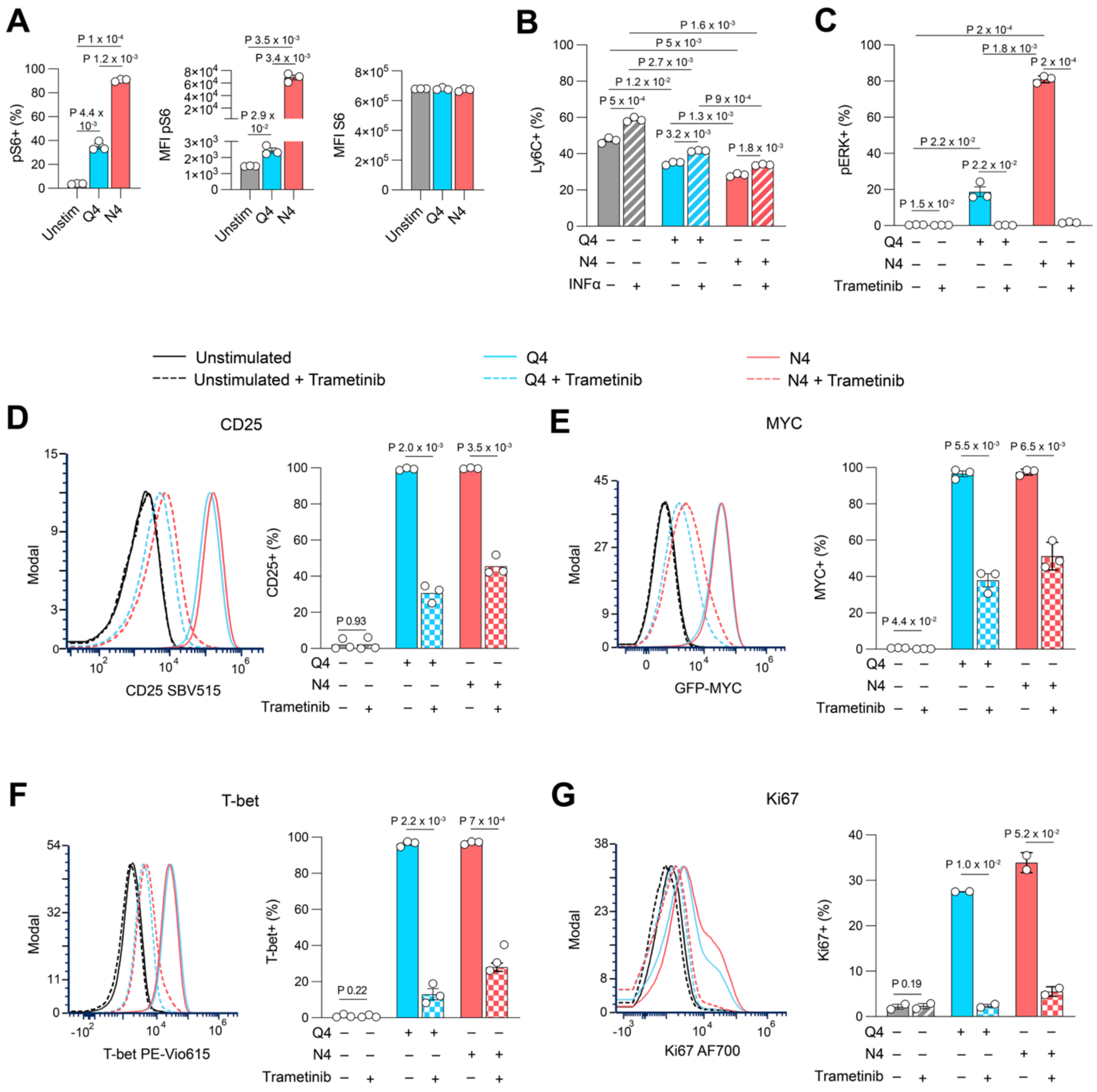

**Supplemental Figure 5. TCR-ligand affinity effects on signaling and impact of ERK1/2 pathway inhibition on *in vitro* activation.** OT-I *Rag2*-/- or GFP-*Myc* OT-I *Rag2*-/- splenocytes were activated *in vitro* with N4 or Q4 peptides, with IFNα (1000 U/ml) or the MEK1/2 inhibitor Trametinib (0.03μM) added where indicated. Cells were assessed by flow cytometry and gated on CD8 T cells. (**A**) pS6 and total S6 measurements following stimulation with N4 or Q4 peptides for 1h. (**B**) Ly6C expression 24 hours post-stimulation. (**C**) Confirmation of Trametinib-mediated inhibition of ERK1/2 phosphorylation after 1h and measurement of effect on (**D**) CD25, (**E**) MYC-GFP, (**F**) T-bet and (**FG**) Ki67 expression after 24h. (**A-C**) Points depict technical replicates, representative of two biological replicates. P-values by Student's t-test. (**D-G**) Bar plots depict combined results from 2-3 biological replicates. P-values by paired t-test, pairing conditions with and without Trametinib within each biological replicate. (**A-G**) Bar height and error bars indicate mean ±SEM.

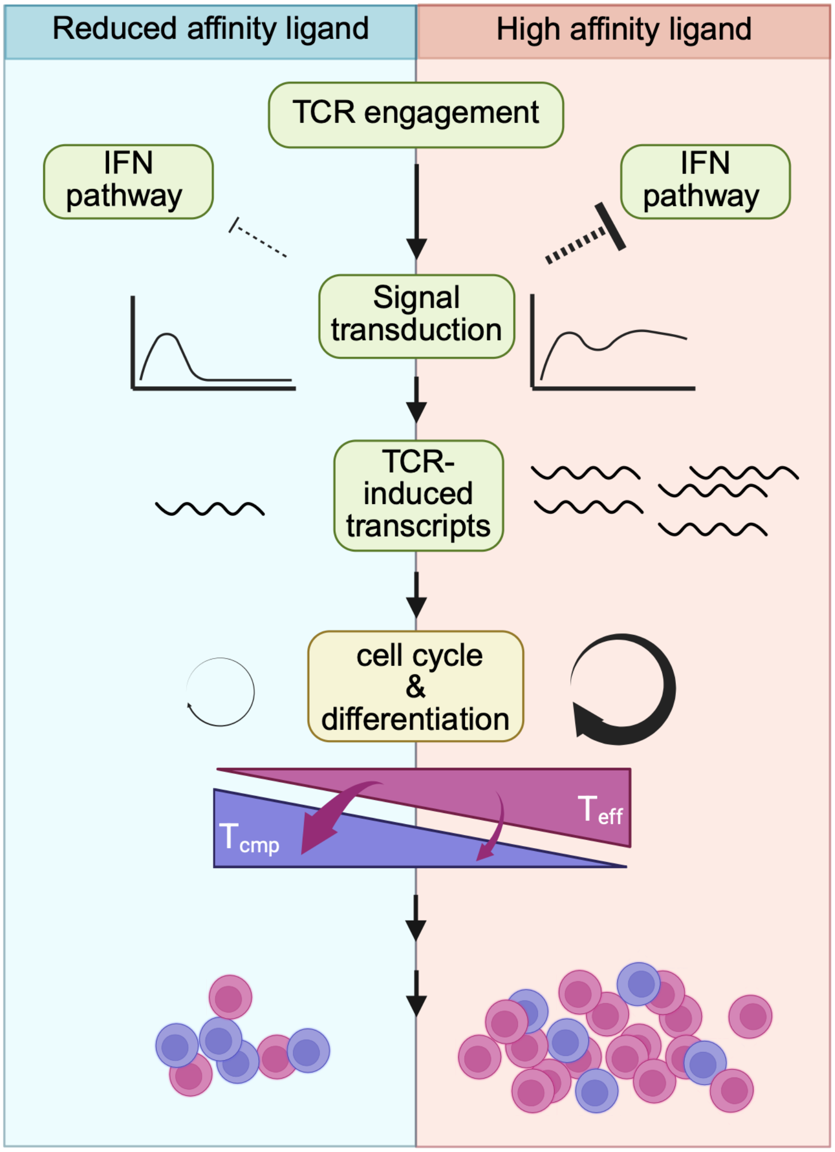


**Supplemental Figure 6. Graphical representation of proposed model connecting TCR-ligand affinity to differentiation fate.**

**Supplemental Tables:**

**Supplemental Table 1.** Cluster-associated marker genes.

**Supplemental Table 2.** Metabolic task enrichment for each cluster.

**Supplemental Table 3.** Differential abundance analysis results comparing cluster membership of N4- versus Q4- derived cells on day 5 post-infection (depicted in Figure 5D).

**Supplemental Table 4.** Pseudobulk differential expression analysis results comparing N4- versus Q4-derived cells within the “acute TCR stimulation” and “initial priming” clusters and N4- and Q4-derived “initial priming” cells to those in the “naïve and weak activation” cluster (depicted in Figures 6A and 7A,C).

**Supplemental Table 5.** Results of Gene Ontology enrichment analysis of differentially expressed genes (depicted in Figure 6C and 7D).

**Supplemental Table 6.** Results of transcription factor binding site enrichment analysis among genes differentially expressed between N4- and Q4-derived cells in the “initial priming” cluster (depicted in Figure 7H).
